# Cross-species single-cell atlases chart progression, therapy-driven remodelling and immune evasion in pancreatic cancer

**DOI:** 10.64898/2026.03.19.712924

**Authors:** Daniele Lucarelli, Shrey Parikh, Sara Jiménez, Christian Schneeweis, Devi Anggraini Ngandiri, Philipp Putze, Tina Kos, Deelaka Wellappili, Vanessa Gölling, Manzila Kuerbanjiang, Caylie Shull, Marie Roja Litwinski, Tania Bori Handschuh, Yasamin Dabiri, Magdalena Zukowska, Barbara Seidler, Raphael Kfuri-Rubens, Stefanie Bärthel, Lennard Halle, Jeanna M. Arbesfeld-Qiu, Dennis Gong, Günter Schneider, Roland Rad, Chiara Falcomatà, Marc Schmidt-Supprian, William L. Hwang, Fabian J. Theis, Dieter Saur

## Abstract

Pancreatic ductal adenocarcinoma (PDAC) is typically diagnosed at advanced stages, yet single-cell datasets that capture late-stage and treated disease remain sparse, hindering progress in understanding tumour heterogeneity and therapy resistance. Here, we have generated integrated single-cell transcriptomic atlases of human and mouse PDAC to define the cellular and molecular landscape of the disease, from early to advanced and metastatic stages, including post-treatment disease, and to enable direct cross-species comparison. Using scANVI to harmonize 16 human studies comprising 257 donors and representative mouse models (101 tumours), we compiled over 1.6 million cells and established a four-level hierarchical taxonomy of more than 60 distinct cell states spanning malignant, stromal, immune, endothelial, adipose, exocrine and endocrine compartments. We resolve ten malignant programmes linked to progression and uncover rare immune phenotypes, including CD4⁺CD8⁺ double-positive T cells that remain poorly characterized in PDAC. Notably, we show that radiotherapy (RT) exposure is associated with enrichment of an EMT-persistent malignant state and an immunosuppressive microenvironment characterized by expansion of tumour-associated endothelium, depletion of intratumoral T cells and heightened laminin–CD44 signalling, with RT-associated genes linked to adverse prognosis in independent cohorts. Cross-species mapping reveals that orthotopic syngeneic allografts more faithfully recapitulate the cellular diversity and EMT-enriched states of advanced human PDAC, underrepresented in autochthonous genetically engineered models, with differences driven primarily by cell-type composition rather than pathway divergence. Together, these atlases and pretrained models provide a broadly accessible reference for benchmarking PDAC model fidelity and for interrogating mechanisms of tumour progression, microenvironmental remodelling and therapy response and resistance.

## Main

Pancreatic ductal adenocarcinoma (PDAC) remains one of the most lethal solid malignancies, with a 10-year survival rate of approximately 1% despite decades of intensive research ^1^. More than 80% of patients present with locally advanced or metastatic disease, and even those undergoing resection almost invariably relapse ^2^. These dismal outcomes are widely attributed to a combination of profound malignant plasticity, a dense fibrotic and immunosuppressive tumour microenvironment (TME) and intrinsic and acquired resistance to systemic and local therapies ^2,3^. Yet, our understanding of how these features emerge, interact and evolve across the clinical course of PDAC, particularly in advanced and treated tumours, remains incompletely understood.

Single-cell RNA sequencing (scRNA-seq) studies have begun to deconstruct PDAC into diverse malignant, stromal and immune cell states, revealing basal-like and classical malignant programs, heterogeneous cancer-associated fibroblast (CAF) subsets and profoundly dysfunctional T-cell compartments ^4^. However, existing datasets are typically limited to early-stage, treatment-naïve resectable tumours from single centres, with modest sample sizes ^4^. As a result, advanced and post-treatment PDAC, the clinical reality for most patients, are underrepresented, and it has been difficult to systematically analyse how therapies such as chemo- or radiotherapy remodel malignant states, vasculature and immunity at a systems level ^4,5^. In parallel, the rapid accumulation of independent scRNA-seq datasets has produced a patchwork of only partially overlapping cell-state taxonomies, making it challenging for the community to compare new datasets, align annotations or confidently place rare populations into a common reference framework ^4^.

Mouse models are indispensable for mechanistic studies and preclinical testing in PDAC, yet their fidelity to human disease at the single-cell level has not been investigated systematically and thus remains poorly defined ^3,4^. Genetically engineered mouse models (GEMMs) recapitulate the initiating key genetic events and aspects of desmoplasia and immune suppression ^3,4,6^, but their capacity to mirror the cellular complexity and therapy-shaped ecosystems of advanced human PDAC is uncertain. Orthotopic syngeneic allograft models are widely used for interventional studies, but the extent to which they capture malignant plasticity, microenvironmental states and treatment-induced phenotypes is largely unknown. A systematic, cross-species mapping between human PDAC and the major classes of mouse models is therefore essential to benchmark model fidelity and to guide model selection and interpretation for specific biological questions and therapeutic interventions.

Addressing these unmet needs requires a unified, cross-species single-cell reference that spans early and advanced disease, incorporates pre- and post-treatment samples, and provides robust, reusable tools for data integration and annotation. Such an atlas should (i) harmonize disparate studies into a shared latent space, (ii) offer a hierarchical and biologically interpretable catalogue of malignant and microenvironmental states, (iii) capture rare but potentially functionally important cell populations, (iv) quantify therapy-associated remodelling of tumour and stromal compartments and (v) allow direct comparison of widely used mouse models with human tumours.

Integration of large, heterogeneous single-cell datasets remains a central challenge in achieving this goal. Atlas-scale efforts must contend with nested batch effects, protocol variability, differences in tissue processing and sequencing chemistry, and inconsistent annotation schemes across studies and platforms. Recent benchmarking and best-practice publications have highlighted that integration methods and pre-processing choices can strongly influence the balance between batch correction and biological signal preservation. They emphasize the need for principled, quantitatively evaluated data integration frameworks ^7,8^. This challenge is further amplified in cross-species analyses, where evolutionary divergence, species-specific gene repertoires and annotation differences complicate the identification of conserved versus species-restricted programs.

Here, we have generated integrated single-cell transcriptomic atlases of human PDAC and mouse models that address these gaps and challenges. Leveraging advances in computational biology and AI-driven harmonization, we apply probabilistic modelling and quantitative benchmarking metrics to align cell states across donors into a shared latent space. The resulting atlases provide a coherent reference for systematic cross-species comparison, capturing the full spectrum of malignant, stromal, immune and endothelial diversity, and delineating conserved programs of tumour progression, microenvironmental remodelling and therapy response. They resolve ten malignant cell states, identify rare immune phenotypes in the PDAC microenvironment, and link prior radiotherapy to an EMT-persistent malignant program and an immunosuppressive, endothelial-expanded, T cell-depleted niche marked by heightened laminin–CD44 signalling and poor-prognosis gene signatures. Together, these atlases and pretrained models provide the community a broadly-accessible, high-fidelity and extensible framework for choosing optimal *in vivo* PDAC models and for elucidating and validating the molecular and cellular mechanisms driving tumour progression, TME and immune remodelling, and response and resistance to therapy.

### The Integrated Core Human PDAC Atlas

To construct a core single-cell transcriptomic atlas of human PDAC, we integrated 11 publicly available sc-and snRNA-seq datasets spanning 191 donors and more than 800,000 cells from primary tumours, adjacent normal pancreas, and metastatic lesions **(Fig. 1, see Methods, Supplementary Table 1)**. Standard highly variable gene (HVG) selection and off-the-shelf integration workflows failed to adequately align tumour and non-tumour compartments across modalities, likely due to the pronounced transcriptional heterogeneity of malignant cells and differences in capture efficiencies between scRNA-seq and snRNA-seq. To overcome these limitations, we implemented a tailored feature selection strategy combining unsupervised Factor Analysis with MOFA ^9^ and expert-guided curation, yielding a biologically coherent set of 2,505 genes used for integration of both species **(see Methods)**. To further minimize technical variation while preserving biological signals, we applied binning on raw counts **(see Methods)** and benchmarked across multiple state-of-the-art integration approaches ^8^. Among all evaluated methods, scANVI ^10^ performed best, providing strong batch correction while maintaining high biological conservation **(Extended Data Fig. 1a, b)**.

**Figure 1.**
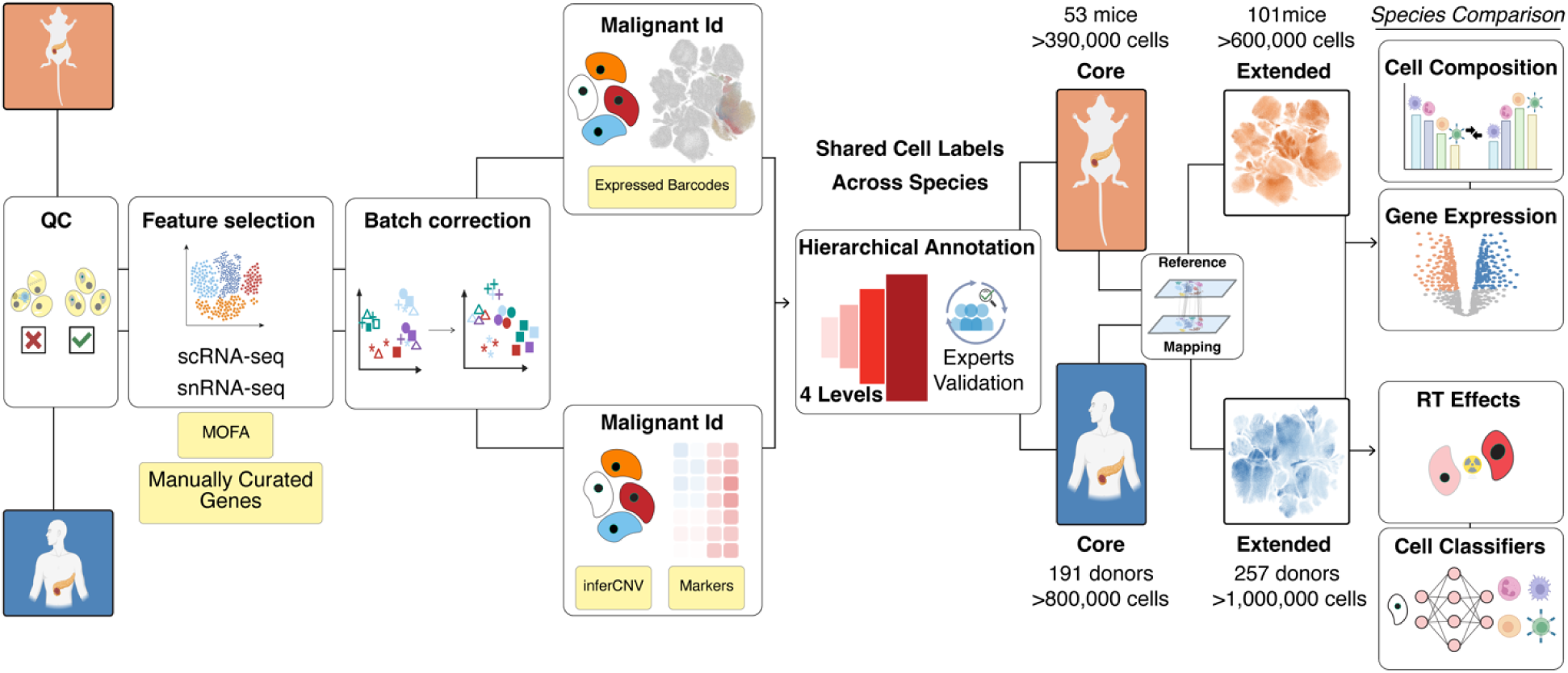
Workflow for constructing translational human–mouse PDAC single-cell atlases. Left: the workflow to build the core atlases and their extension, including (i) quality control (QC); (ii) feature selection combining literature-derived and expert-curated markers with unsupervised matrix factorization using MOFA ^9^; (iii) benchmarking of seven batch correction methods; (iv) malignant cell annotation in mouse through expressed barcodes; and (v) a four-level hierarchical annotation externally validated by PDAC experts. Middle: the human core was built from single-cell/single-nucleus RNA sequencing count data from 191 donors, while the mouse core atlas was generated from 53 individual tumours. Each core atlas was then extended through reference mapping to additional datasets, yielding extended atlases covering 257 human donors and 101 mouse samples. For the extended atlases, reference mapping of new datasets was performed onto the respective core atlas (human-to-human, mouse-to-mouse). Right: examples of downstream applications: the extended atlases were used to model effects of radiotherapy (RT), train cell-type classifiers for community use, and to enable systematic cross-species comparisons.

The integrated atlas resolved the PDAC ecosystem into three major compartments: epithelial, immune, and stromal cells (Level-1 annotation), which are further divided into seven major lineages at Level-2: malignant, lymphoid, stromal, myeloid, exocrine, endothelial and endocrine cells **(Fig. 2a)**. This multi-lineage framework provides the foundation for a four-level hierarchical annotation comprising more than 60 distinct cell states **(Fig. 2a-f, Extended Data Fig. 2a-g)**. These levels range from broad lineages to progressively fine cellular subtypes, and culminate in detailed annotations of context-dependent cell states, which represent functional transcriptional programs underlying phenotypic heterogeneity within a given cell type **(Fig. 2 and Extended Data Fig. 2)**. Initial malignant versus non-malignant cell assignment was based on copy number variation profiles inferred with inferCNV (**see Methods),** while canonical lineage markers from previous studies were used to establish broad identities. We then applied expert-curated and validated molecular signature sets **(Supplementary Table 2-3),** which were used to assign cells to distinct states by systematic scoring and thresholding **(Extended Data Fig. 2a-g)**. Where marker overlap introduced ambiguity - for example, between EMT (epithelial-mesenchymal transition) and mesenchymal malignant states - we applied a k-nearest neighbour classifier trained on the latent space obtained through scANVI^10^ to refine boundaries and confidently annotate cells **(see Methods)**. Finally, annotations were independently reviewed and validated by domain experts, who were provided with the cell clusters and their corresponding marker gene expression profiles blinded to our prior cell-state assignments. Each expert independently assigned cell states based on marker expression, and a majority-vote consensus across experts defined the final cell state labels **(see Methods)**.

**Figure 2.**
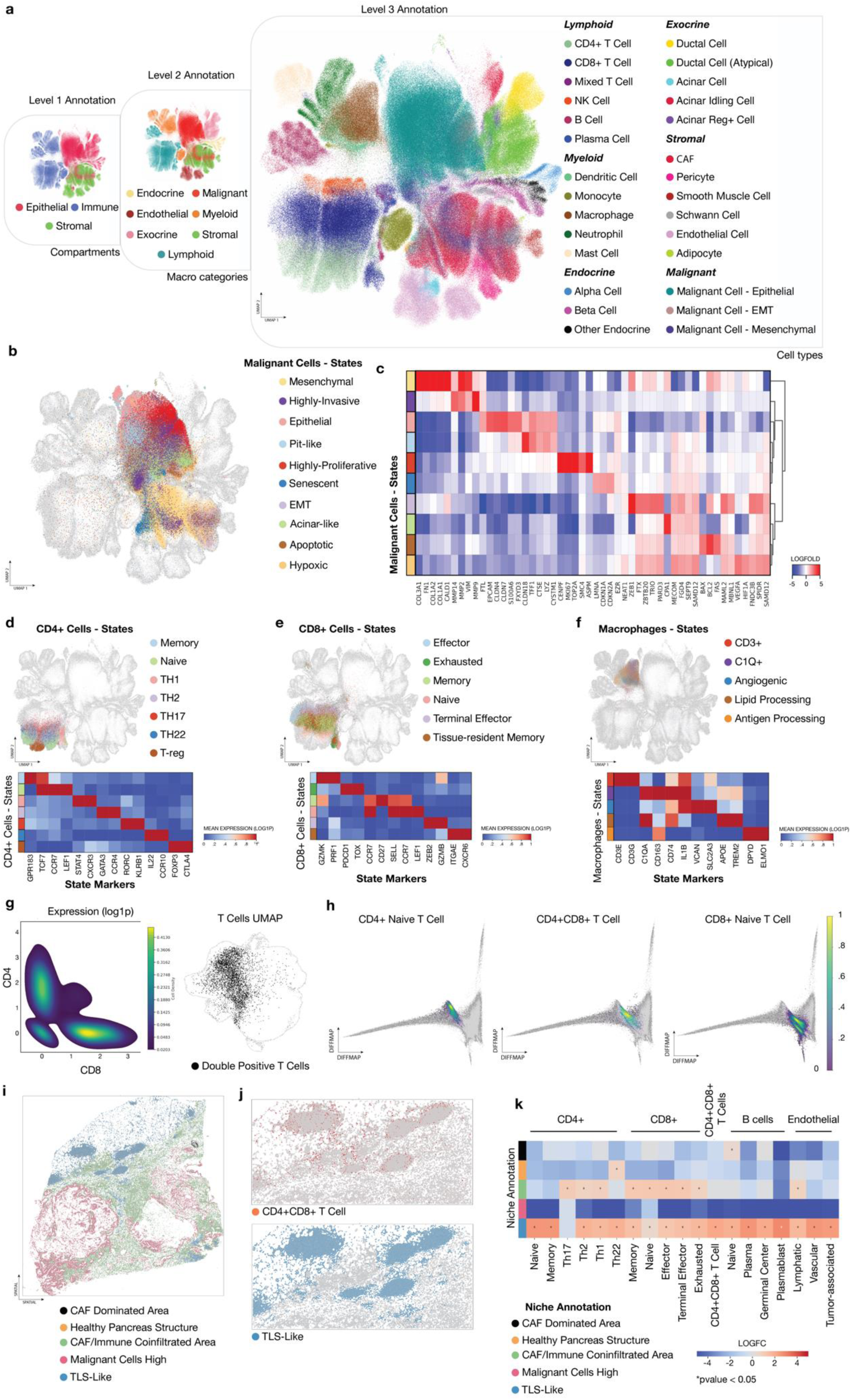
The integrated PDAC single-cell atlas enables hierarchical and fine cell-state annotation in the Human Core Atlas. **a)** UMAP of the integrated human PDAC atlas showing hierarchical annotation at three levels — Level 1 (major compartments), Level 2 (macro categories), and Level 3 (fine-grained cell-type resolution). **b)** Subclustering of malignant cells (Level 4) resolves ten transcriptionally distinct states: epithelial, EMT, mesenchymal, hypoxic, highly-proliferative, highly-invasive, senescent, apoptotic, pit-like and acinar-like. **c)** Heatmap of representative marker genes delineating each malignant cell state. **d-e)** CD4⁺ and CD8⁺ T cell subtypes and associated markers identified within the lymphoid compartment, including naïve, memory, helper, effector, exhausted, terminal-effector, tissue-resident and regulatory (T-reg) populations. **f)** Macrophage subclustering reveals distinct macrophage cell states, including C1Q⁺, angiogenic, lipid-processing, CD3⁺ TAMs and antigen-presenting populations. **g)** Validation of DP T cells using density plots of canonical markers (CD4/CD8) and UMAP projections highlighting the double-positive T cell population. **h)** Density map of T cell subsets on diffusion map manifold showing naïve CD4+ (left), DP CD4+ CD8+ T cells (middle) and naïve CD8+ (right) localization. **i)** Spatial transcriptomics (Xenium) PDAC section from one patient ^19^ segmented into recurrent tissue niches: CAF dominated area, healthy pancreas structure, CAF-immune co-infiltrated area, malignant cells high, and tertiary lymphoid structure (TLS)-like. **j)** Spatial maps from the same section showing inferred localization of CD4⁺CD8⁺ DP T cells in red (top) and TLS-like in blue (bottom). **k)** Differential enrichment of immune and endothelial subsets across indicated spatial niches of six patient samples profiled by spatial transcriptomics ^19^, shown as log fold-change (logFC), grouped by lineage (CD4⁺, CD8⁺, CD4⁺CD8⁺ DP T cells, B cells, endothelial).

Within the malignant compartment, we resolved ten transcriptionally distinct states that extend beyond the classical-basal dichotomy: Epithelial, EMT, Mesenchymal, Hypoxic, Highly-proliferative, Highly-invasive, Senescent, Apoptotic, Pit-like and Acinar-like **(Fig. 2b and Extended Figure 2b)**. These states reflect the heterogeneity of PDAC cells, their diverse adaptation strategies, and capture key functional transcriptional programs. For example, hypoxic cells were enriched for *HIF1A* and *VEGFA*, epithelial cells for *EPCAM* and *KRT8/18*, mesenchymal cells for collagens, highly-invasive cells for *MMP2/9/14*, and Pit-like cells harbour a foveolar/pit-epithelial gene signature ^11^, including *GKN1, GKN2*, and *CLDN18* **(Fig. 2c and Extended Data Fig. 2b)**. Non-malignant epithelial populations, including acinar and ductal cells, as well as multiple stromal lineages, such as adipocytes, CAFs and endothelial cells were similarly resolved at higher resolution **(Extended Data Fig. 2a, c, d)**. The immune compartment displayed marked cellular diversity, encompassing multiple T cell subsets, B cell and NK cell populations **(Fig. 2d-e, Extended Data Fig. 2e-f)**, and a broad spectrum of myeloid cells, including various tumour-associated macrophage (TAM), monocytes, neutrophil, and dendritic cell subtypes **(Fig. 2f, Extended Data Fig. 2g)**.

The depth and breadth of the integrated atlas enabled robust identification of cell populations that had been rarely detected or only anecdotally reported, but now emerge consistently across donors and datasets. These include CD4^+^CD8^+^ double-positive T cells and CD3^+^ macrophages, previously described only in isolated reports and other settings ^12–16^ **(Fig. 2g-h, Extended Data Fig. 2e,g,h)**. Differential expression analysis of CD4⁺CD8⁺ double-positive (DP) T cells revealed co-expression of canonical DP markers (CD4, CD8B) together with a distinct transcriptional signature **(Extended Data Fig. 2e, Supplementary Table 4)** enriched for naive/central memory-associated genes (*IL7R*, *CCR7*, *TCF7*, *LEF1*), redox regulators (*TXNIP*, *SESN3*), and mitochondrial metabolic genes (*ATP5*, *LDHB*), consistent with a transitional or stress-adapted naïve/central memory-like state within the PDAC TME, which is distinct from the activated cytotoxic DP T cell program reported in other tumours ^12,13,17,18^.

Diffusion map analysis supported this interpretation as DP T cells occupied an intermediate position along the diffusion manifold between CD4⁺ naive and CD8⁺ naive T cell states **(Fig. 2h)**, consistent with a transitional, early-differentiation phenotype rather than a terminal effector program. This placement suggests that DP cells may reflect a shared naive-/central memory-like compartment bridging the CD4⁺ and CD8⁺ naive continua, potentially arising from lineage plasticity or transient co-receptor expression. To further validate the existence of DP cells and investigate their spatial localization in PDAC, we leveraged an independent spatial transcriptomics PDAC dataset ^19^ and transferred our atlas-derived cell-state annotations onto the spatial profiles **(see Methods)**. We then used CellCharter ^20^ to infer spatial cellular niches, resolving five main regions: CAF dominated area, healthy pancreas structures, CAF/Immune co-infiltrated area, malignant cell high areas, and tertiary lymphoid-like structures (TLS-like), based on the cell type compositions in the niches **(Fig. 2i)**. Strikingly, DP T cells were preferentially localized at TLS-associated niches, rather than broadly distributed across immune-infiltrated tumour regions contrary to more differentiated T cells **(Fig. 2j-k)**. This spatial enrichment supports the diffusion-based placement of DP cells along a naive-/central memory-like continuum and demonstrates that DP T cells are linked to TLS-like niches or lymphoid-structured regions within the PDAC TME.

CD3^+^ TAMs expressed canonical macrophage markers alongside a complete T-cell receptor (TCR) complex (*CD3D/E/G*, *TRAC*, *TRBC1/2*) and effector genes (*GZMA*, *CCL5*, *NKG7*) **(Supplementary Table 4)**. This population exhibited mixed myeloid and T-cell transcriptional programs suggestive of lineage infidelity, or fusion events ^21,22^ **(Extended Data Fig. 2g; Supplementary Table 4)**. Similar dual-identity macrophages have been reported in other solid tumours ^16,21,22^ and may contribute to immune modulation or evasion in PDAC.

Collectively, this integrated core atlas delivers the most comprehensive single-cell reference of human PDAC to date. By harmonizing data across diverse studies and platforms, it delineates the cellular and transcriptional architecture of the PDAC TME, captures the full spectrum of malignant and stromal diversity, and reveals previously underappreciated hybrid and transitional cell states that provide a foundation for mechanistic and translational follow-up.

### Extension of the Core Human PDAC Atlas

To demonstrate the utility and scalability of the core atlas, we extended it by incorporating 5 additional datasets comprising 66 donors and over 270,000 cells. After count binning using the same gene set as for the core atlas integration, the additional query datasets were mapped into the existing latent space via the pretrained scANVI ^10^ model using the scArches framework ^23^. This transfer-learning approach allowed seamless incorporation of new data without full reintegration of all datasets, demonstrating the scalability and adaptability of our atlas framework **(Fig. 3a)**.

**Figure 3.**
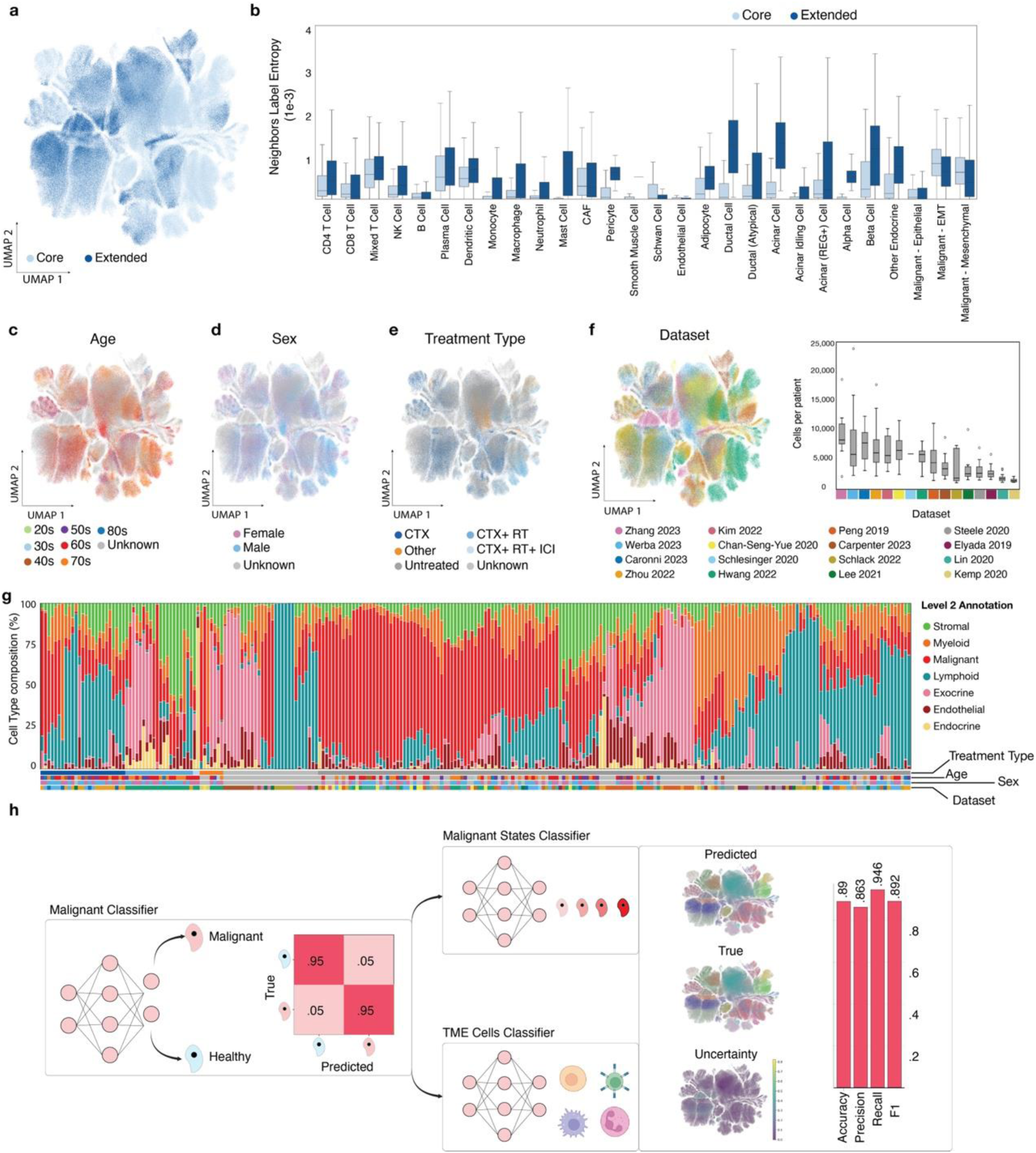
The extended human PDAC atlas retains cross-study coherence and supports accurate classification of malignant and TME cell states. **a)** UMAP of the integrated human PDAC atlas showing the core datasets (11 studies) and the 5 additional datasets included in the extension. **b)** Neighbourhood label entropy across cell types showing consistent integration quality and preservation of lineage structure between core and extended datasets. **c-f)** Distribution of cells in the integrated atlas by age (c), sex (d), treatment type (e), and dataset of origin (f), illustrating broad demographic and experimental diversity. The boxplot in panel f shows per-dataset cell count distributions. **g)** Cell-type composition across samples, coloured by Level 2 annotations and stratified by metadata (sex, age, treatment type, dataset), highlighting diverse tumour–stroma–immune proportions across cohorts. **h)** Schematic of the hierarchical classification framework built on the atlas: A first-stage multi-layer perceptron (MLP) network distinguishes malignant from non-malignant cells (left panel), followed by specialized classifiers for malignant and TME cell types/states (Level 3). Right panels: UMAP projections comparing predicted versus true labels, uncertainty estimates and classifier performance (accuracy, precision, recall and F1 score). CTX, chemotherapy; RT, radiotherapy; ICI, immune checkpoint inhibition; F1, harmonic mean of precision and recall metrics.

To quantitatively assess mapping fidelity between the core and extended datasets, we compared the neighbourhood label entropy of cells originating from either the core or the extension, computed using a weighted Shannon entropy score **(see Methods)**, based on Level 3 annotations **(Fig. 3b)**. Core cells showed uniformly low entropy, while extension cells exhibited only slightly higher entropy, consistent with the expected uncertainty when projecting unseen data. The small median difference of < 0.3 × 10⁻³ across cell types indicated that the overall structure of major cell lineages is preserved. Importantly, common malignant and immune populations retained low entropy values (∼0.5–1.0 × 10⁻³), demonstrating robust and consistent transfer of fine-grained annotations. In contrast, higher entropy (2–3 × 10⁻³) was observed mainly in normal ductal and acinar compartments, likely reflecting transcriptional overlap with malignant epithelial states rather than annotation noise. Together, these results establish the PDAC atlas as a stable and extensible reference that can integrate new datasets while preserving cell-type resolution.

The extended atlas now comprises 16 studies, 257 donors and over one million cells, providing the research community a rich resource to explore PDAC heterogeneity across a broad spectrum of patient cohorts and therapeutic conditions **(Fig. 3c-f, Supplementary Table 5).** Higher level compositional analyses revealed a stratification of patients at the macro cell type level, dividing them into branches with high proportion of malignant and stromal cells characterized by low immune infiltration, and, conversely, patients with low percentage of malignant cells and high immune infiltration, whether from the myeloid or lymphoid branch (**Fig. 3g**).

Having established the robustness and scalability of the PDAC human atlas with extensively annotated Level 4 cell states **(Extended Data Fig. 3, Supplementary Table 3)**, we next built a hierarchical classification framework to enable the PDAC research community to rapidly transfer fine-grained atlas annotations to new PDAC scRNA-seq datasets without explicitly mapping them to the atlas (**Fig. 3h**). We trained a multi-layer perceptron (MLP)–based classifier in two stages: a first-stage model separates malignant from non-malignant cells, followed by specialised MLPs that assign cells to Level 3 atlas categories within the malignant and TME compartments (**see Methods)**. The malignant vs non-malignant classifier achieved >95% correct prediction, and the specialised classifiers showed high accuracy, precision and recall across cell states (**Fig. 3h**). In summary, the extended PDAC atlas and associated classifiers provide a robust foundation for scalable data integration and reproducible, atlas-level annotation, enabling community-wide interrogation of PDAC heterogeneity at single-cell resolution.

### Radiotherapy is Associated with Enrichment of Stable EMT Malignant States, TME Remodelling and T Cell Exclusion

Therapeutic interventions, including chemotherapy and radiotherapy (RT), are known to remodel the PDAC TME and influence malignant cell-state dynamics; however, large-scale single-cell comparisons across these interventions remain limited ^3–5,24^. To systematically assess the impact of current standard-of-care therapies, we leveraged the analytical power of our PDAC atlas and first examined variation in malignant cell-state composition across cohorts. Unsupervised clustering of patient-level malignant cell-states composition revealed five robust subgroups characterized by distinct malignant programs: C1 (Epithelial), C2 (Early Transition), C3 (EMT-Intermediate), C4a (Mesenchymal), and C4b (EMT-Persistent) **(Fig. 4a)**. Pseudotime trajectories positioned C1→C4a along a canonical epithelial-to-mesenchymal continuum (C1→C2→C3→C4a), consistent with progressive dedifferentiation and acquisition of mesenchymal features. Notably, a second trajectory diverged from C3 toward C4b, which displayed a stable, EMT-enriched transcriptional program and appeared to represent a persistent EMT endpoint rather than a transient intermediate state **(Fig. 4a)**.

**Figure 4.**
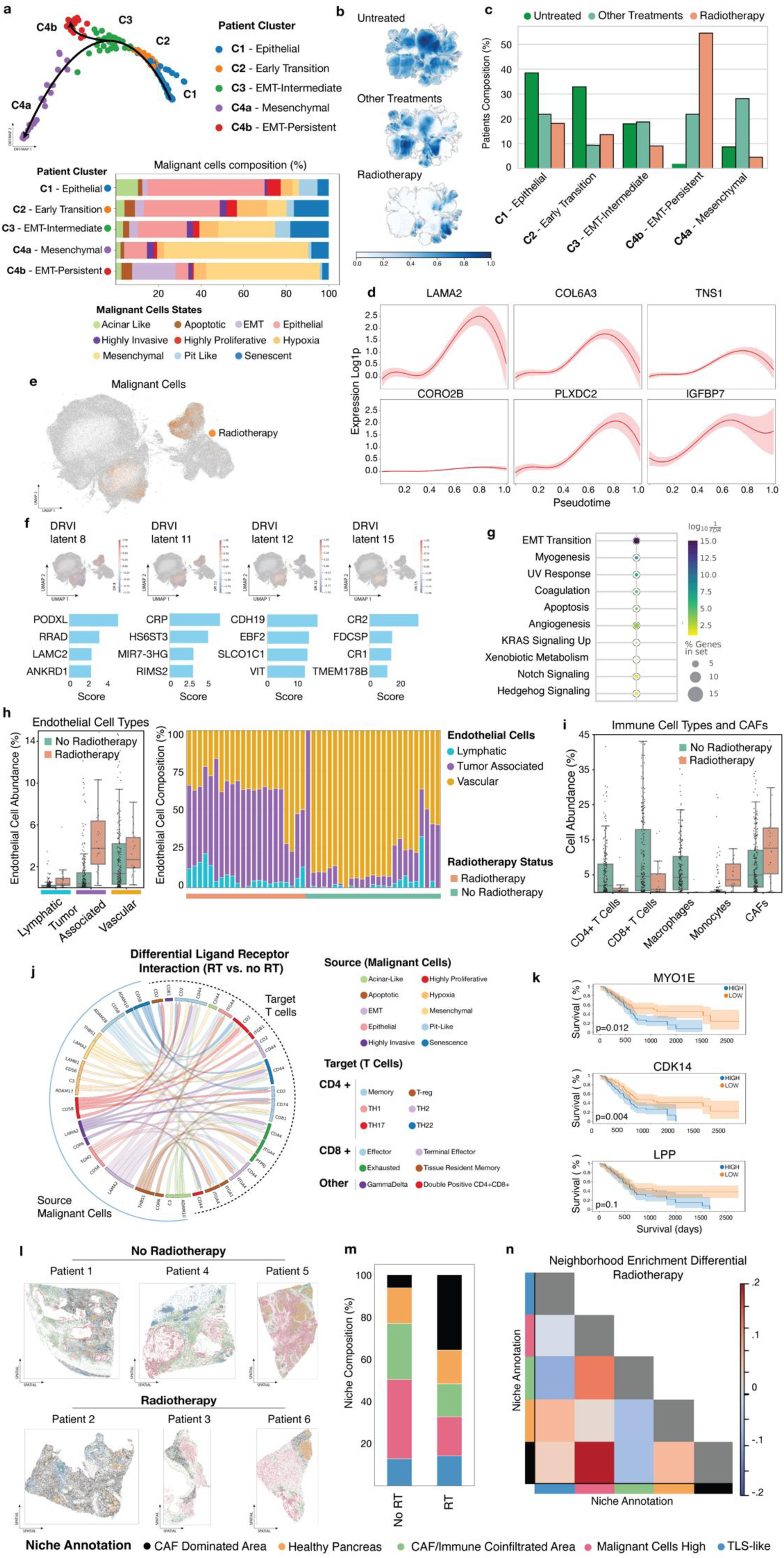
Radiotherapy exposure drives a terminal EMT-state, endothelial expansion, and immune exclusion in PDAC. **a**) Diffusion Map with patient-level composition manifold. Each point represents one patient as a 10-dimensional vector of malignant state proportions (percent of the ten Level-4 malignant states). Pseudotime and clustering on this manifold were computed on this composition space, revealing a progression from Epithelial (C1) → Early Transition (C2) → EMT-Intermediate (C3) → Mesenchymal (C4a), and a distinct EMT-Persistent terminal endpoint (C4b). Stacked bars below show malignant-state composition over patient clusters. **b)** UMAPs of patients based on therapeutic intervention (upper panel, untreated; middle panel, other treatments (combinations of different treatments without radiotherapy exposure); lower panel, radiotherapy. **c)** Distribution of treatment modalities (untreated, radiotherapy, and combinations of different treatments without radiotherapy exposure) across patient clusters, showing a marked enrichment of RT-treated samples within EMT-Persistent clusters (C4b). **d)** Gene expression dynamics (pseudotime) along the Epithelial → EMT-Persistent trajectory, highlighting progressive activation of extracellular-matrix and stromal-activation programs, chemokines/stromal recruitment factors, complement/immune modulators and transcriptional regulators. **e)** UMAP of malignant cells integrated with the interpretable DRVI model ^28^ coloured by RT exposure, demonstrating segregation of RT-treated cells. **f)** Identification of interpretable latent factors using Deep Disentangled DRVI. Here, DR (e.g., DR8, DR11) denotes specific disentangled latent factors that capture distinct underlying gene programs enriched in RT-treated tumour cells. **g)** GSEA of the genes defining these RT-enriched disentangled factors, showing enrichment in EMT transition, myogenesis, angiogenesis, apoptosis, KRAS-, Hedgehog-, and Notch-signalling. **h)** Endothelial cell composition across RT groups, demonstrating expansion of lymphatic and tumour-associated vascular endothelium in RT-treated tumours (left) and tumour-associated endothelium being the major responsible cell subset (right). **i**) Boxplots showing depletion of multiple CD4⁺ and CD8⁺ T-cell subsets and an increase of CAFs in RT-treated tumours, consistent with immunosuppressed highly vascularized microenvironments. **j)** Differential ligand-receptor cell-cell communication analysis between RT and non-RT treated tumours inferred with LIANA+, summarizing top RT-enriched interactions between malignant states and lymphoid populations, enriched for ECM–integrin and laminin–CD44 axes. **k)** Kaplan–Meier survival analysis of PDAC patients in TCGA-PAAD for differentially expressed RT-associated genes (*MYO1E*, *CDK14, LPP*), illustrating that high expression of adhesion/EMT regulators correlates with reduced overall survival (Log-rank test, p values indicated in the individual panels) **l)** Spatial transcriptomics PDAC sections of RT treated (bottom) and non-treated (top) tumours ^19^ revealed five recurrent tissue niches: CAF dominated, healthy exocrine, CAF/immune co-infiltration, malignant cells high, and tertiary lymphoid structure like (TLS-like). **m)** Group-level niche composition comparing no RT and RT tumours, displayed as stacked bars of niche fractions, highlighting increased CAF infiltration territory and reduced malignant cells high regions in RT-exposed samples. **n)** Differential neighbourhood enrichment analysis between niches in RT versus no RT tumours. Heatmap shows logfold changes in niche–niche spatial co-localization (RT relative to no RT); red indicates increased neighbourhood enrichment under RT and blue indicates decreased enrichment. CAF, cancer-associated fibroblasts; RT, radiotherapy; DR, disentangled component of DRVI latent space.

We next linked these malignant programs to therapeutic interventions. At the compositional level, RT-exposed tumours not only exhibited the highest proportion of EMT malignant cells, but also represented the dominant fraction of the total composition of the C4b EMT-Persistent cluster **(Fig. 4a-c)**. To define transcriptional programs underlying this shift, we conducted a pseudobulk trajectory-driver analysis of malignant cells at the patient level using pseudotime ^25^ kernel with CellRank ^26,27^, complemented by an interpretable single-cell transcriptomic integration of RT- and non-RT-treated malignant cells using DRVI ^28^. Trajectory driver analysis along the C3 to C4b (EMT-persistent) axis revealed enrichment of cell-matrix adhesion and cytoskeletal regulators (e.g., *LAMA2*, *COL6A3*, *TNS1*, *CORO2B*), consistent with reinforcement of an ECM/adhesion-driven invasive phenotype **(Fig. 4d)**. DRVI-based latent factor analysis further identified genes upregulated in RT-treated malignant cells, including *LAMC2*, *PODXL*, *TGFB1*, *ANGPTL4*, *RRAD*, *MEOX2*, *SLCO1C1*, *PI16*, *APOD* and *CDH19* (**Fig. 4e,f**), with gene-set enrichment profiles implicating EMT transition, angiogenesis, myogenesis, KRAS-, Notch- and Hedgehog-signalling **(Fig. 4g)**.

Consistent with these malignant cell programs, RT exposure was also associated with a substantial remodelling of the TME. Compositional analyses revealed expansion of tumour-associated vascular and lymphatic endothelial cell subsets in RT-treated tumours **(Fig. 4h)**. In parallel, immune and CAF profiling revealed intratumoral T cell depletion in RT patients, affecting both CD4+ and CD8+ subsets, as well as an increase in CAF abundance **(Fig. 4i)**, compatible with a more vascularized yet immunosuppressed, T cell excluded desmoplastic TME.

To dissect the underlying communication routes, we performed differential ligand-receptor analysis with LIANA ^29^ in RT- and non-RT treated tumours separately, followed by ranking of the interactions with the highest delta scores between the two groups focusing on interactions involving malignant cells as source and both endothelial and T cell compartments as target cells **(Fig. 4j, Supplementary Table 6 and 7)**. In RT-exposed samples, malignant-to-lymphoid signalling was dominated by enhanced engagement of the laminin-CD44 axis (LAMA2/LAMB1->CD44) across multiple T-cell subsets; an interaction previously implicated in limiting T cell chemotaxis and tissue infiltration ^30^. Additionally, we observed increased ADAM10-CD44 engagement in RT-treated tumours, consistent with enhanced sheddase activity targeting surface CD44 that may further impair T cell trafficking and retention ^31–33^. Notably, RT has been shown previously to induce ADAM10 and fibrosis in PDAC models ^34^. Accordingly, pharmacologic or genetic targeting of ADAM10 in mice decreased RT-induced fibrosis and tissue tension, reduced tumour cell migration and invasion, sensitized orthotopic tumours to radiation killing, and prolonged mouse survival ^34^.

To evaluate clinical relevance, we evaluated RT-associated differential gene expression changes in malignant cells for prognostic associations in the TCGA-PAAD cohort using Cox proportional hazards models **(Fig. 4k)**. *MYO1E* and *CDK14*, both implicated in cytoskeletal dynamics and cell–matrix interactions, were significantly associated with reduced overall survival (log-rank p = 0.012 and 0.004, respectively; HRs ≈ 1.4–1.3). *LPP* showed a similar, though non-significant trend toward poorer outcomes (p = 0.10; HR = 1.46). These results align with the observed enrichment of EMT- and ECM-related programs in RT-treated tumours and support a model in which radiation exposure selects for invasive, adhesion-driven malignant states embedded in a vascularized, T cell excluded niche that is associated with adverse prognosis.

To validate our findings, we analysed spatial transcriptomics data of RT-treated patient tissue specimens ^19^. RT-treated samples showed a pronounced expansion of CAF-dominated niches **(Fig. 4l)**, indicating a shift toward desmoplasia, alongside a contraction of immune infiltrates within the immune-/CAF-dominated regions **(Fig. 4m)**. We next quantified how RT altered niche-to-niche spatial organization using differential neighbourhood enrichment. This analysis indicated that RT restructures local tissue architecture by increasing the proximity of malignant cell high regions to CAFs and TLS neighbourhoods **(Fig. 4n).** Collectively, these spatial analyses reinforce a model in which RT drives the reorganization of the PDAC microenvironment, characterized by redistribution of malignant foci into stromal-dominated neighbourhoods and concomitant immune exclusion.

### The Integrated Mouse PDAC Atlas

Although single-cell transcriptomic studies have transformed our understanding of PDAC, translational frameworks that directly link human disease states to the preclinical models used to study them are still missing ^4^. Existing large-scale atlases catalogue cellular diversity but generally do not quantify how faithfully different model systems capture key phenotypes observed in patients. To close this gap, we constructed an integrated Mouse PDAC Atlas using the same hierarchical framework as the human atlas, thereby enabling direct cross-species comparison and quantitative benchmarking of model fidelity.

A central challenge in annotating PDAC at single-cell resolution is the transcriptional similarity between fibroblasts and mesenchymal malignant cells, which can confound malignant-stromal boundaries. To establish an unambiguous ground truth for malignant identity, we used expressed barcodes ^35^ to label cancer cells prior to orthotopic implantation into syngeneic, immunocompetent mice, allowing barcode-based separation of tumour cells from host-derived stroma and immune populations in the core atlas (**Fig. 5a**). Additional non-barcoded in-house and public single-cell datasets were then mapped onto this embedding following a similar extension workflow as established for the human atlas. The resulting final extended mouse PDAC atlas comprises over 600,000 cells from 101 individual tumours, spanning a diverse set of samples originating from genetically engineered mouse models (GEMMs) and orthotopic, syngeneic allografts in immunocompetent hosts (**Fig. 5b**). We included available metadata, such as sex, treatment and model type, and observed, as expected, partial segregation by model, indicating that some orthotopic and endogenous autochthonous GEMMs represent biologically distinct ecosystems rather than interchangeable experimental surrogates (**Fig. 5b-e, Supplementary Table 8-9**).

**Figure 5.**
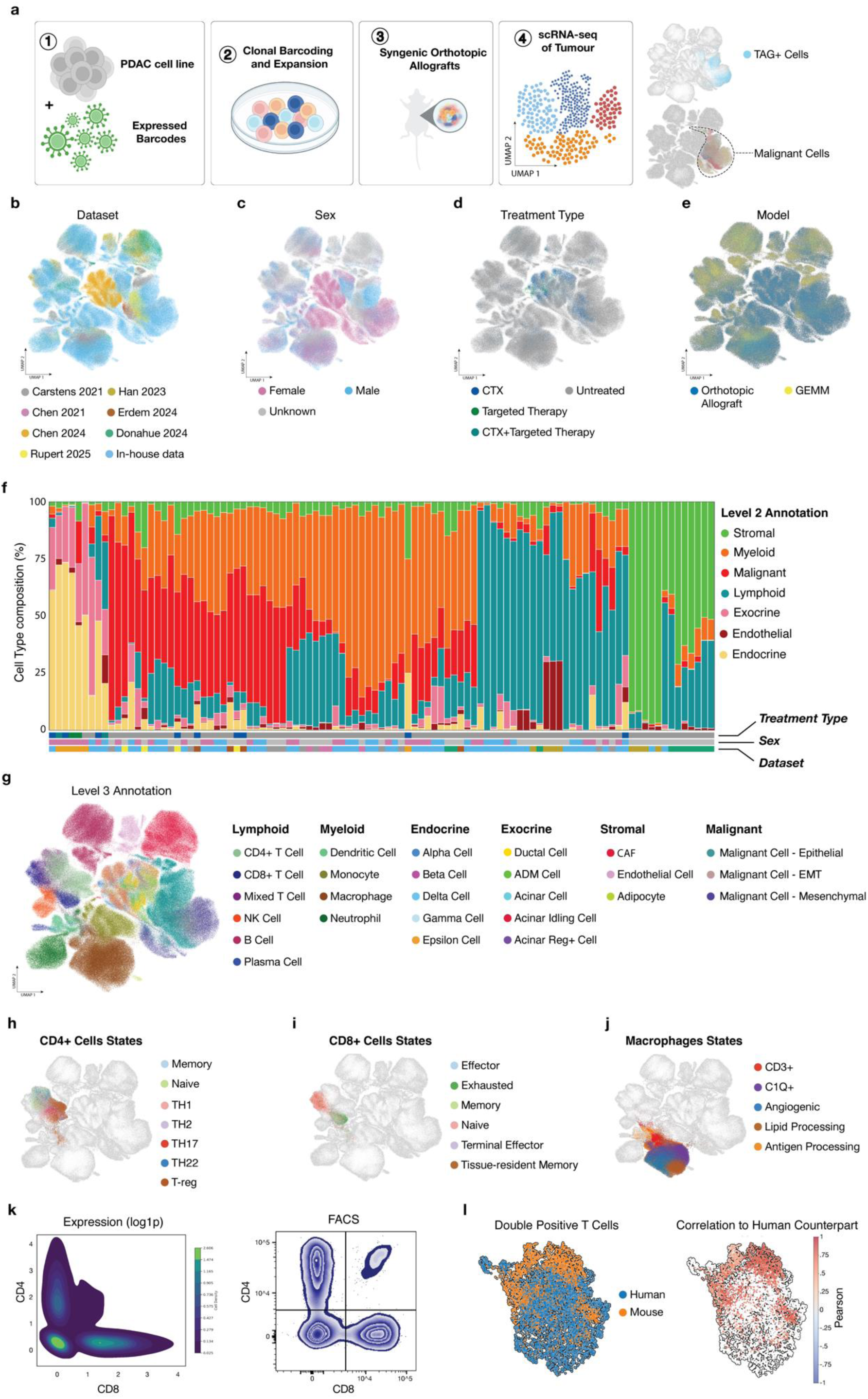
Construction of an integrated Mouse PDAC Atlas with a harmonized cell-type hierarchy mirrors the Human PDAC Atlas for cross-species comparison. **a)** Schematic overview of the experimental design. Mouse PDAC cell lines were clonally barcoded using a lentiviral library (1), expanded and orthotopically transplanted into syngeneic immunocompetent mice (2–3), followed by scRNA-seq profiling of resultant tumours (4). Expressed barcode tracing enabled unambiguous separation of malignant (TAG⁺) from host-derived non-malignant cells (right panel, UMAPs depicting barcoded (TAG^+^) (top) and malignant (bottom) cells). **b-e)** UMAPs of the integrated Mouse PDAC Atlas coloured by dataset (b), sex (c), treatment type (d), and model (orthotopic syngeneic immunocompetent allografts vs. autochthonous GEMMs (e). **f)** Sample-wise cell-type composition across treatment types, datasets, and models. Autochthonous tumours were dominated by classical epithelial-like malignant states, whereas orthotopic allografts displayed greater heterogeneity with an enrichment of EMT, hypoxic, and mesenchymal programs. **g**) Level 3 hierarchical annotation of the Mouse Atlas using the same multi-tiered scheme as the Human Atlas, resolving lymphoid, myeloid, stromal, endocrine, exocrine, endothelial, and malignant compartments. **h-j)** Substate resolution of major immune and stromal lineages: CD4⁺ T cells (h), CD8⁺ T cells (i), and macrophages (j), showing distinct regulatory, effector, angiogenic, and lipid-processing programs. **k)** Validation of double-positive (DP) CD4⁺CD8⁺ T cells at the transcriptomics level (transcription density plots, left panel) and at the protein level by flow cytometry (right panel). **l)** UMAP showing DP T cells coloured by species (left) and Pearson correlation of mouse DP T cell gene expression against the human DP T cell archetype (right).

To enable direct cross-species comparisons, we maintained the same hierarchical annotation framework as in the human atlas and resolved the mouse atlas into the seven major Level 2 cellular lineages: myeloid, lymphoid, malignant, stromal, exocrine, endocrine, and endothelial. Compositional analysis revealed extensive heterogeneity in the relative representation of different compartments across samples, mirroring the inter-patient heterogeneity seen in human PDAC (**Fig. 5f)**. Within each lineage, cells were further annotated using the same marker-based criteria as in the human atlas, yielding overlapping Level 3 cell-types and Level 4 cell-states across species **(Fig. 5g-j, Extended Data Fig. 4a-g and 5)**. This unified labelling framework supports consistent cross-species mapping, analysis, and modelling. Using this framework, we identified analogous CD4⁺ and CD8⁺ T-cell states, as well as distinct macrophage polarizations, closely mirroring those in the human atlas **(Fig. 5h–j)**.

We next asked whether the unconventional double-positive CD4⁺CD8⁺ T cell and CD68⁺CD3ε⁺ macrophage subpopulations detected in the human atlas are also present in murine tumours. We examined the corresponding mouse populations and observed subsets of cells co-expressing CD4/CD8 and CD68/CD3ε at the transcript level, and confirmed their existence at the protein level by flow cytometry, validating these double-positive populations across species **(Fig. 5k, Extended Data Fig. 4e,g,h)**. To investigate cross-species similarity of double positive (DP) T cells, we integrated human and mouse DP T cells using a shared latent space learned with scVI ^36^ and visualized with UMAP **(Fig. 5l, left panel)**. For each mouse DP T cell, we then computed the Pearson correlation between its expression profile and the human DP T cell meta cell profile across the whole orthologous gene space, revealing strong conservation of DP T cells and their associated immune states across species **(Fig. 5l, right panel)**. Together, these analyses indicate that DP immune states observed in human PDAC have conserved transcriptional counterparts in mice and provide a quantitative framework for cross-species benchmarking and mechanistic interrogation.

Overall, the mouse PDAC atlas establishes a directly comparable reference to the human atlas, positioning preclinical models within the broader landscape of human PDAC. Built within a unified hierarchical framework and anchored by barcode-defined ground truth for malignant cells, it enables systematic cross-species analyses, reveals model-specific strengths and limitations, and contextualizes human disease states in experimentally tractable systems.

### Cross-Species Mapping Reveals Conserved Malignant Programs and Model-Specific Cellular Composition in PDAC

To systematically assess conservation of transcriptional programs and tissue organization across species, we performed a cross-species comparison of the Human and Mouse PDAC atlases. This analysis enabled direct comparison of malignant cell states and microenvironmental composition across autochthonous GEMMs, syngeneic orthotopic allografts, and human patient samples within a single reference framework. We first computed presence scores ^37^ as a proxy for the likelihood that specific cell states identified in humans are represented in the mouse atlas, stratified by model type. GEMMs were enriched for epithelial states, whereas orthotopic tumours displayed a marked increase in proliferative, mesenchymal, apoptotic and EMT-like programs **(Fig. 6a)**. These patterns indicate closer alignment of orthotopic models with advanced-stage human PDAC, which is characterized by a higher burden of mesenchymal tumour cells.

**Figure 6.**
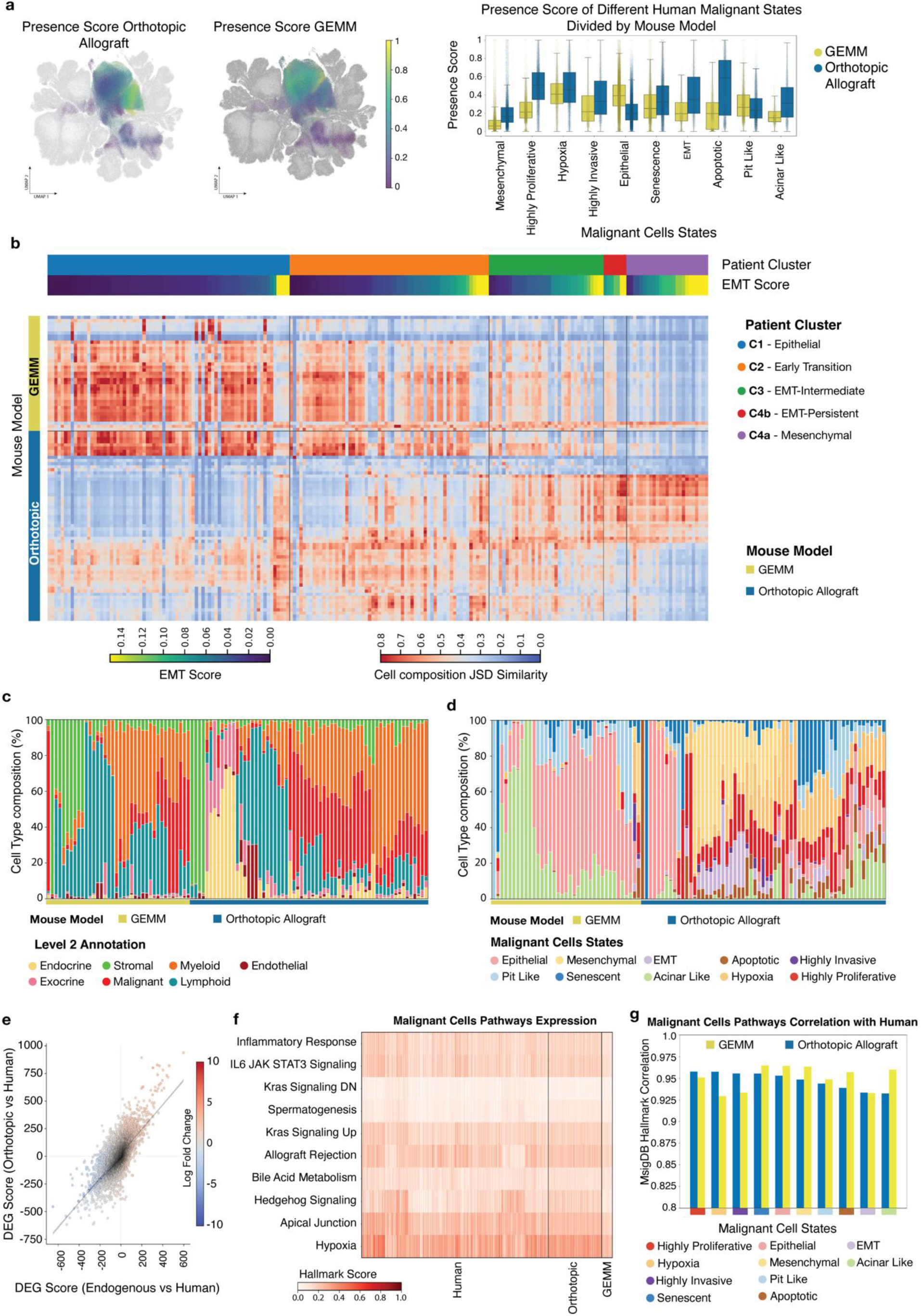
Cross-species comparison of the Mouse and Human PDAC atlases reveals conserved malignant-state programs and microenvironmental organization. **a)** Presence scores showing the probability of human malignant cell states to be found in the Mouse PDAC Atlas. Left UMAP depicts orthotopic allografts; right UMAP depicts autochthonous GEMMs. Scores are projected onto the human UMAPs (left panel), and stratified by malignant cell state (right panel - bar plots). Epithelial malignant states have higher presence scores in autochthonous GEMMs, whereas highly proliferative, mesenchymal, apoptotic, and EMT-like programs have a higher presence score in the orthotopic allografts indicating increased transcriptional plasticity and closer resemblance to advanced human PDAC. **b)** Cross-species compositional overlap between mouse and human PDAC samples. Similarity computed as the inverse of Jensen–Shannon divergence (JSD) of cell type distributions between human patient clusters (as defined in Fig. 4) and mouse models. Orthotopic tumours display greater overlap with late-stage or EMT-enriched human clusters, whereas endogenous models align more with epithelial-like human states. **c-d)** Sample-wise compositional profiles across all cell types (c, Level 2 annotation) and malignant cell states (d, Level 4 annotation), stratified by model type. Orthotopic tumours show a higher proportion of malignant cells overall and a pronounced shift toward EMT, hypoxia, and mesenchymal states, while endogenous GEMMs retain more differentiated epithelial profiles. **e)** Differentially expressed gene (DEG) concordance between mouse and human malignant cells. Both, orthotopic allografts (y-axis) and autochthonous GEMMs (x-axis) show high correspondence across models. **f)** Expression levels of selected cancer hallmark pathways (from MSigDB) show high overlap between the orthotopic and autochthonous GEMM models and the human counterpart. **g)** Cross-species correlation of malignant-cell pathway enrichment profiles, demonstrating similar signalling patterns between models and high correlation with the human PDAC counterpart, further supporting the conservation of malignant-state programs across species.

To quantify global cross-species similarity, we computed a composition-based similarity score defined as the inverse of the Jensen–Shannon divergence (JSD) between mouse and human cellular profiles. Orthotopic allografted tumours displayed significantly higher similarity to late-stage and EMT-enriched human clusters, while autochthonous GEMMs more closely resembled epithelial-dominant early-stage human tumours **(Fig. 6b)**. Together, these findings support the notion that orthotopic tumours recapitulate the molecular and cellular heterogeneity of clinically advanced human PDAC.

We next examined cellular composition across tumour models in more detail. At the global level, orthotopic tumours contained a higher proportion of malignant cells **(Fig. 6c)**. Within the malignant compartment, autochthonous tumours were dominated by epithelial populations, whereas orthotopic tumours were enriched for mesenchymal and EMT-associated states **(Fig. 6d)**, in line with the broader transcriptional shifts observed above. These results suggest that, in orthotopic settings, malignant and microenvironmental compartments co-evolve towards greater plasticity and aggressiveness, mirroring advanced human PDAC, while GEMMs predominantly model earlier, more epithelial stages.

To dissect transcriptional conservation beyond composition, we compared malignant cell states across the models and their human counterparts. We performed state-resolved differential expression analysis using the human malignant states as a reference framework. For each state, we computed differentially expressed genes (DEGs) separately for orthotopic vs human, and autochthonous vs human comparisons, and correlated the resulting log fold-change vectors. Both comparisons revealed strong concordance, indicating that core malignant transcriptional programs are preserved across mouse models and species **(Fig. 6e)**. We then performed gene enrichment analysis on these state-specific signatures and correlated pathway enrichment profiles between orthotopic, autochthonous and human tumours. This showed high overlap in the enriched pathways between different malignant cell states across the three systems **(Fig. 6f-g, Supplementary Table 10)**. Collectively, these analyses demonstrate that malignant cells share remarkably conserved gene expression and signalling programs across human tumours, orthotopic allografts and GEMMs. While orthotopic tumours more closely mirror the cellular complexity and transcriptional features of advanced human disease, the differences between allografts and autochthonous GEMMs arise primarily from shifts in tumour composition and malignant cell states abundances rather than divergence of the underlying malignant gene expression patterns **(Fig. 6c-g)**. This cross-species atlas thus provides a principled basis for selecting models tailored to early epithelial versus advanced EMT-rich PDAC biology.

## Discussion

We present comprehensive single-cell transcriptomic atlases of human and mouse PDAC that establish a unified molecular framework for interpreting tumour heterogeneity, therapy-associated remodelling and cross-species similarities. Through separate integration of over 1.6 million single cells from 257 human PDAC and 101 mouse samples, our study provides a harmonized and hierarchically annotated reference that resolves PDAC into major cellular lineages and over sixty fine-grained states spanning malignant, stromal, endothelial, and immune compartments. The human PDAC atlas represents, to our knowledge, the most extensive integration of PDAC single-cell datasets to date and enables robust comparison across studies and platforms. The hierarchical annotation framework captures a wide spectrum of PDAC cell states - from broad lineages to specialized transcriptional states - providing a stable reference for reproducible annotation and downstream analysis.

Beyond providing a unified reference for human–mouse PDAC ecosystems, our integrative analysis uncovered rare and unconventional immune populations, including CD4⁺CD8⁺ double-positive (DP) T cells and CD3⁺ tumour-associated macrophage-like cells co-expressing T-cell receptor (TCR) components, which have so far been underrecognized and poorly characterized ^12–16,21,22,38^. Although historically often dismissed as technical artifacts, their recurrence across independent datasets and species, as well as across different cancer entities ^12,13,15,16^ suggests that they represent bona fide components of the PDAC microenvironment. While CD3⁺ macrophages remain poorly understood ^15,16,21,22^, DP T cells have recently been described in several cancer types as tumour-infiltrating lymphocytes whose differentiation and function can be shaped by local antigenic stimulation and microenvironmental cues ^12,13^.

Across immunotherapy-responsive cancer types, DP T cells can adopt at least two non-exclusive functional modes: (i) tumour-reactive cytotoxic effectors enriched in immune checkpoint blockade responders ^12,13^; and (ii) helper or immunoregulatory states, including Th2-skewed cytokine production and reduced cytotoxicity ^39,40^. These observations suggest DP T cells are plastic and an extrathymic source of single-positive T cells whose net impact likely depends on the tumour type, tissue niche, antigen exposure, and local circuits ^12,13,39^. In PDAC, we observe a strikingly different DP phenotype. Trajectory analyses position them between naïve CD4⁺ and naïve CD8⁺ states, consistent with a naïve/central-memory–like program marked by low cytotoxicity/exhaustion and enrichment of lymphoid-homing features ^12,17,18^. Spatially, they are found predominantly within TLS-like regions and are infrequent outside these niches. This spatial restriction is notable and links DP T cells to PDAC TLS biology, which has been associated with an immune-active microenvironment and improved clinical outcome, with higher T and B cell infiltration and reduced immunosuppressive infiltrates in TLS⁺ tumours ^41,42^. In addition, TLS can function as organized immune microanatomical sites: in PDAC patients they can be induced, for example, by GM-CSF-secreting whole-cell pancreatic cancer vaccine (GVAX) approaches ^43,44^.

These findings support a model in which TLS-associated DP T cells may constitute a spatially restricted reserve compartment, poised for activation and differentiation, yet held in a quiescent, multipotent state. This idea resonates with the emerging concept that productive antitumor immunity is sustained by lymphoid-like niches that preserve less-differentiated T cell states capable of proliferative renewal ^45,46^. Future studies will need to determine their antigen specificity, differentiation potential, and relationship to TLS maturation and local inhibitory pathways ^38,41,43,44^.

The atlas was designed as an extensible resource that can be incrementally expanded as new datasets emerge. By applying scArches-based transfer learning, we demonstrated that new studies can be seamlessly incorporated without full re-integration, preserving annotation consistency and biological structure ^23^. The pretrained scANVI model and associated hierarchical classifier enable rapid mapping of external single-cell datasets onto the reference, offering an accessible and powerful tool for the PDAC research community ^10^. Importantly, extensive benchmarking across integration strategies confirmed the technical robustness of this framework. Compared with previous PDAC atlases, which vary in depth and lineage resolution ^47,48^, our hierarchical annotation provides a comprehensive and standardized definition of PDAC cell states and supports exploration of malignant, stromal, and immune diversity at unprecedented granularity.

This unified framework also enables systematic investigation of how therapy reshapes PDAC ecosystems. By integrating clinical metadata, we observed that RT is associated with pronounced remodelling of both, the malignant and the surrounding microenvironment. RT-treated tumours were enriched for EMT and mesenchymal-like malignant cell states and displayed an expansion of tumour-associated endothelial lineages ^24,49^, together with a marked reduction of multiple intratumoral CD4 and CD8 T cell subsets. These findings are consistent with clinical and translational studies reporting that chemoradiation can profoundly alter the PDAC immune microenvironment, often reducing cytotoxic T-cell infiltration and increasing suppressive and regulatory populations ^50–52^. Latent factor and cell-cell interaction analyses highlighted extracellular matrix (ECM) remodelling, integrin and cytoskeletal signalling, angiogenesis, and Notch/TGFb-associated EMT programs as dominant RT-associated malignant states. Within this context, we identify a distinct EMT-persistent malignant endpoint that branches from the canonical epithelial-to-mesenchymal continuum and is particularly enriched in RT-exposed tumours, extending preclinical work that linked RT to EMT induction, invasive behaviour and metastasis in PDAC and other cancer types ^34,53^. Concurrently, our interaction analyses point to mechanistic axes that may couple these malignant adaptations to immune exclusion. In RT-exposed tumours, malignant-to-lymphoid signalling is dominated by laminin–CD44 (LAMA2/LAMB1→CD44) and ADAM10–CD44 interactions, in line with functional work implicating laminin–CD44 circuits and ADAM10-mediated CD44 shedding in regulating lymphocyte adhesion, migration and tissue retention ^30–34,54,55^. These data support a model in which RT fosters adhesion- and ECM-driven malignant states that reside in laminin-rich, T cell excluded niches. The identification of RT-associated adhesion and cytoskeletal genes such as *MYO1E* and *CDK14* as markers of adverse prognosis in independent cohorts links this transcriptional remodelling to clinical outcome and suggests that at least part of the detrimental effect of RT in PDAC may be mediated by selection for EMT-persistent, ECM-anchored, immune-excluded clones. These observations align with, and extend, prior reports of RT-induced fibrosis, desmoplasia and therapy resistance in PDAC and other cancer types ^34,52,53,56–58^, and raise the possibility that rational combinations targeting, for example, ECM–integrin signalling, ADAM10 activity, or laminin–CD44 interactions may be required to fully exploit RT in this disease ^50,59–63^.

To bridge preclinical and human biology, we constructed a first-of-its-kind mouse PDAC atlas encompassing over 600,000 cells from genetically engineered and orthotopic allografted tumour models using the same hierarchical annotation. This enabled direct cross-species comparisons of malignant and microenvironmental states. Strikingly, orthotopically transplanted syngeneic allografts exhibited greater compositional and transcriptional similarity to advanced human PDAC, characterized by mesenchymal and EMT-enriched malignant states, whereas autochthonous genetically engineered models (such as KPC) were predominantly dominated by epithelial-like states more reminiscent of earlier-stage disease. Despite these compositional differences, core malignant transcriptional programs, including EMT, hypoxia, proliferation, and RAS effector signalling, were highly conserved between human and mouse, indicating that fundamental PDAC circuitry is preserved across species and that model differences arise largely from shifts in cell-type abundance and state distribution rather than pathway-level divergence ^64^. The identification of analogous CD4⁺CD8⁺ DP T cells and CD3⁺ macrophages in both human and mouse PDAC further underscores this conservation ^12,15^ and provides an entry point for mechanistic interrogation of these dual-identity populations in tractable model systems.

Together, our findings establish a cross-species reference for PDAC that unifies the molecular, cellular, and therapeutic dimensions of the disease. The atlas provides a foundation for systematic evaluation of model fidelity, facilitates annotation and integration of future datasets, and enables discovery of conserved and context-specific programs that drive PDAC progression and treatment resistance. Several limitations should be acknowledged. The current integration includes limited representation of metastatic lesions and healthy pancreas, restricting the ability to reconstruct full disease trajectories from normal tissue to precursor lesions and metastatic states. Future efforts combining single-cell, single-nucleus and spatial transcriptomics, alongside multi-omic and cross-species integration, will be instrumental in overcoming these limitations.

To ensure broad utility of this work, we will make the complete human and mouse PDAC atlases - integrated matrices, metadata, hierarchical annotations, together with pretrained scANVI and classifier models - openly accessible. The pretrained models enable researchers to project their own single-cell datasets directly into the atlas latent space using the scArches framework, allowing rapid and reproducible assignment of malignant and microenvironmental cell states and enabling direct comparison of new datasets against the atlases without requiring advanced computational expertise. Detailed tutorials and code examples will accompany the release, providing guidance on dataset mapping, hierarchical state annotation label transfer, and cross-species comparison. The atlases will be made available through web platforms enabling investigators to contextualize new data within a unified reference framework. By releasing both the integrated atlases and pretrained models, we aim to provide an evolving community resource that not only accelerates mechanistic insights into pancreatic cancer biology but also informs the design of novel therapies, including rational combinations of stromal- and immune-targeted interventions.

## Methods

### Collection and Preprocessing of Human Datasets

We collected 11 publicly available single-cell and single-nucleus RNA sequencing (scRNA-seq and snRNA-seq) datasets from Gene Expression Omnibus (GEO), ensuring the inclusion of metadata from associated manuscripts (**Supplementary table 1**). For pre-processing, we applied consistent quality control across all datasets. To remove potential outliers, cells with total counts below the 5th percentile or above the 95th percentile were excluded. Genes expressed in less than 10 cells were removed, as well as cells expressing fewer than 400 genes. This stringent preprocessing ensured high-quality data for downstream analysis. Furthermore, we collected 5 additional publicly available studies for the extension of the core atlas. The preprocessing was done in a similar fashion to the core atlas.

### Data Normalization

Raw count matrices from each dataset were normalized using Scanpy’s normalize_total function (target sum = 1 × 10⁶) followed by logarithmic transformation (log1p).

### Annotation

Annotation of cell types was performed in a hierarchical manner. For scRNA-seq data, Leiden clustering was initially performed at a resolution of 0.2, and differentially expressed (DE) gene analysis was conducted to identify coarse annotations, classifying cells into broad categories: immune, stromal, epithelial/malignant, endothelial, erythroid, neuronal, and endocrine. Ambiguous clusters were refined with clustering at a lower resolution of 0.1. Subclusters within each Leiden cluster were further identified at a resolution of 0.2, and DE analysis was conducted to assign Level 1 annotations with more granularity. This multi-level annotation approach facilitated detailed characterization of cell populations across datasets.

### Copy Number Variation Analysis and Malignant Cell Annotation

To infer large-scale copy number variation (CNV) patterns indicative of malignancy, we applied inferCNV (via the infercnvpy package; https://github.com/broadinstitute/infercnv) to each dataset individually. Individual datasets were used as batch_covariate, and genes expressed in fewer than five cells were removed. For each dataset, epithelial populations (annotated as Acinar cell, Ductal cell, or Ductal/Malignant cell) were designated as potentially malignant, whereas all other cell types served as reference “normal” populations. CNV inference was performed with a genomic window size of 150 genes using the default parameters. The resulting CNV expression matrices were subjected to principal component analysis (PCA), neighbourhood graph construction, Leiden clustering, and UMAP embedding. CNV scores were computed for each cell, providing a quantitative estimate of genomic instability. Cells displaying elevated CNV scores were classified as malignant. Cut-off thresholds for CNV score–based annotations were determined manually for each dataset by visual inspection of score distributions and chromosome heatmaps.

### Binning

Binning was performed using a custom Python function that divides each gene’s non-zero expression values into 50 quantiles. For each cell, non-zero gene counts were digitized into discrete integer categories according to quantile thresholds, with zero values retained as a separate category. This procedure preserves the rank information of expression profiles while minimizing the influence of outliers. The resulting binned data were stored in the adata.layers["binned_data"] field, and corresponding bin thresholds were saved in adata.obsm["bin_edges"] for reproducibility.

### Feature Selection for Integration

A systematic feature selection strategy was employed to identify genes for integration. Initially, 12000 HVGs were identified for scRNA-seq and snRNA-seq datasets individually, of which 6082 were overlapping. MOFA ^9^ was applied on this restricted overlapping features space using batch covariates based on the dataset, resulting in 15 latent factors. From these, 2134 genes were selected by choosing the top 500 genes from top 10 relevant factors after removing overlaps. Additional DE genes per cell type, cell type markers from the Broad Institute, and custom panel markers were incorporated, yielding a final list of 2520 genes, 2505 of which were present in overlapping HVGs. This comprehensive selection ensured robust integration while preserving biologically relevant variation.

### Integration

Integration of scRNA-seq and snRNA-seq datasets was carried out using both supervised and unsupervised probabilistic frameworks to balance biological fidelity and batch correction performance. The integrated feature space was constructed from the final set of high-confidence manually selected genes using the binned expression layer described above. We benchmarked multiple integration methods—scVI ^36^, scANVI ^8^, scPoli ^65^, sysVI ^66^, DRVI ^28^, and expimap ^67^ with default model parameters, treating donor identity as a batch covariate. Among the tested methods, scANVI demonstrated optimal performance according to scIB metrics ^8^, achieving robust alignment across donors and modalities while preserving cell-type–specific expression structure. For visualization, UMAP embeddings were computed using 100 nearest neighbours, a cosine distance metric, and a minimum distance parameter of 0.75 to capture both global and local transcriptional relationships.

### Post Integration Quality Control

Final quality control of the integrated dataset included median absolute deviation analysis to identify and remove outliers per Dataset and per cell type lineage, defined as exceeding three median absolute deviations for metrics such as log-transformed total counts, number of genes per count, and mitochondrial gene percentages ^68^. Empty droplets were identified based on log-normalized MALAT1 expression ^69^, using modality-specific thresholds (scRNA-seq > 3.5, others ≥ 4.0), and removed prior to downstream analysis. In total, approximately 30,000 outlier cells were removed further.

### Re-annotation

The integrated atlas was re-annotated with the guidance of leading experts in the field to ensure biological accuracy and relevance. Hierarchical annotation was refined using clustering-based approaches and validated with known canonical markers specific to PDAC up to Level 3 of annotation. For the finest level annotation of the cell states at Level 4, we further scored the cells with expanded expert curated marker lists, defined thresholds and assigned categorical labels. This expert-driven process ensured robust classification of cell types and subtypes, enhancing the biological interpretability of the dataset and its alignment with current knowledge in pancreatic ductal adenocarcinoma research.

### Domain Expert Annotation Validation

To independently validate cell type annotations, we engaged eight external experts in pancreatic cancer biology who were not involved in the initial cell annotation process. Each expert received de-identified clustering results along with cluster-level gene expression summaries and marker gene enrichments, without access to the original annotations or dataset identities (blinded review). Experts were asked to assign each cluster to one of the predefined major cell categories and further to specify finer subtypes based on the provided gene expression profiles and canonical markers. Their individual classifications were aggregated, and final cluster identities were determined by majority voting across all eight experts. This expert validation ensured robust, unbiased assignment of cellular identities and served as an independent verification of the computationally derived annotations.

### KNN Classifier to Resolve Ambiguous Cell Annotation

To resolve ambiguity amongst cell states of a particular cell type that might be too similar to two or more states, we used a k-nearest neighbours (knn) classifier. The knn classifier was trained with 25 neighbours and on the scANVI latent space on cells already robustly classified on the previous steps. The classifier was then used to predict the label of misclassified cells. The final prediction was added to the Level 4 of annotation.

### Label Entropy Score

To quantify the local heterogeneity of cell states in the k-nearest neighbour (kNN) graph of the extended atlas, we computed a weighted label Shannon entropy for each cell. For a given cell i, the entropy H_i was defined as:

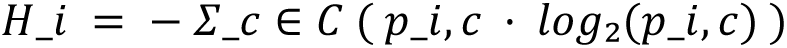

where C is the set of all possible cell-type labels, and p_i,c represents the weighted probability that a neighbouring cell of *i* has label *c*. The probabilities were computed as:

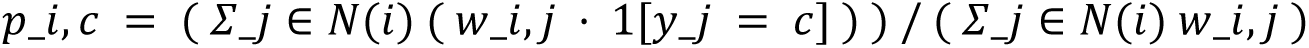

where N(i) denotes the set of neighbouring cells of cell i in the kNN graph, y_j is the label of neighbour j, and w_{i,j} is a distance-based weight between cell i and neighbour j. Specifically, w_{i,j} corresponds to the Euclidean distance d_{i,j} between cells i and j, such that closer neighbours contribute more strongly to the local label composition. This weighted approach accounts for the fact that cells located near transition zones in the neighbourhood graph between two cell states are more likely to have mixed neighbourhoods, while cells within homogeneous regions exhibit lower entropy. Entropy values close to zero indicate that nearly all neighbours share the same label (high neighbourhood purity), whereas larger values indicate locally mixed neighbourhoods.

### Multi-Layer Perceptron for Cell Type Classification

We trained a series of multi-layer perceptron (MLP) classifiers to automate hierarchical cell annotation within the integrated PDAC atlas. The first classifier distinguished malignant versus non-malignant cells using log-normalized gene expression as input features and expert-validated cell type labels as ground truth. The model comprised three fully connected layers (ReLU activation, dropout = 0.2) and was optimized using Adam with categorical cross-entropy loss. Model performance was evaluated on a held-out validation set, achieving 95% accuracy and balanced precision/recall across classes. Predicted cells were subsequently passed to two specialized MLP models: a TME classifier to resolve non-malignant cells and a malignant-state classifier to assign malignant cell type. Both models used identical architectures and training protocols, with label uncertainty estimated via softmax entropy. Model predictions were projected onto the atlas embedding for visualization.

### Patient Stratification, Pseudobulking and Cell-Rank Driver Analysis

We stratified patients by malignant-state composition by constructing a patient-level compositional matrix, where each observation captured the relative abundance of ten malignant cell states. Principal component analysis (PCA), followed by k-nearest neighbour (KNN) graph construction and Leiden clustering, identified five robust patient groups with distinct malignant-state profiles. A diffusion-based pseudotime analysis was applied on this embedding to reconstruct transitions among patient states.

To model transcriptional dynamics along the epithelial-to-mesenchymal (Epi→Mes) axis, we generated pseudobulk objects per patient by aggregating the mean expression per gene, transferred the compositional embedding, and applied CellRank to infer lineage probabilities and nominate trajectory drivers. Transition matrices were computed from diffusion pseudotime kernels, and macrostates were defined using Generalized Perron cluster analysis. Genes exhibiting strong correlation with lineage absorption probability and pseudotime were designated as putative trajectory drivers of malignant-state progression.

### Explainable Integration of Malignant Cells with DRVI

We subset the extended human PDAC atlas to malignant cells from RT-exposed (chemoradiation) and non-exposed (chemotherapy-only) donors, excluding untreated and treatment-unknown cases. To obtain an interpretable low-dimensional representation and identify genes associated with treatment-related variation, we applied Deep Disentangled Variational Inference (DRVI), using donor identity as a batch covariate. Integration was performed using default model parameters. We then identified latent factors that showed strong association with radiotherapy exposure and extracted their top contributing genes. These gene sets were subsequently subjected to gene set enrichment analysis (GSEA) to characterize the biological programs underlying radiotherapy-associated malignant states ^70^.

### Cell-Cell Communication with LIANA

To examine how RT reshapes intercellular signalling, we inferred ligand–receptor (LR) interactions using LIANA (Dimitrov et al. 2024). The extended atlas was subset to RT-exposed and non-exposed tumours, and LIANA’s rank aggregate was run independently on each group using the consensus resource. Analyses were performed on log-normalized expression values grouped by Level 4 refined cell types (expression proportion > 0.1). Results were merged across conditions and summarized by a composite delta-score combining standardized differences in interaction magnitude, probability, and log fold-change (delta = 1.0 × z⍰ₐg + 0.7 × z⍰ᵣₒᵦ + 0.5 × z⍰ₒgFC). Positive delta values indicate preferential engagement in RT. Significant LR pairs (p < 0.05) were ranked within key axes including tumour–endothelial, tumour/CAF–endothelial, tumour/CAF–tumour, and tumour–T-cell interactions, highlighting pathways most rewired by RT.

### Survival analysis

Overall survival analyses were performed using TCGA clinical and transcriptomic data aggregated at the patient level. Survival time was defined as the maximum recorded number of days to death, or otherwise to last follow-up for censored cases. Patients without survival time information were excluded. Multivariable survival analyses were performed using Cox proportional hazards regression with gene expression modelled as continuous variables. Genes with zero variance were excluded prior to modelling. Cox models were fitted using a penalized partial likelihood (L2 penalization, penalizer = 0.1) and adjusted for sex, with hazard ratios and 95% confidence intervals estimated from fitted models. Kaplan–Meier survival curves were generated for visualization by segregating patients into high and low expression groups using the median expression value as the cutoff. Kaplan–Meier analyses were unadjusted and compared using two-sided log-rank tests. All analyses were performed in Python using the lifelines package.

### Presence Score

Similar to HNOCA (He et al. 2024), we quantified how strongly each single cell in the human atlas is represented by mouse tumour models using a weighted cross-species nearest-neighbour framework followed by diffusion smoothing, producing a per-cell presence score for each model (i.e., orthotopic allografts, autochthonous GEMMs). scANVI latent space was used for the analysis separately for malignant and non-malignant compartments segregated by Level 3 annotations. For a given compartment, we first built a mouse→human k-nearest neighbour graph with NNDescent (k = 1000), keeping only query→reference edges. Edge weights captured local neighbourhood concordance using an overlap of human kNN sets (k = 1000). The number of shared neighbours between a mouse cell and a human cell was converted to a Jaccard index and squared to emphasize strong matches while damping weak overlaps. Mouse cells were then aggregated by model: for each model, we summed its mouse→human edge weights across all of that model’s mouse cells to obtain an initial coverage vector over human cells. To denoise and propagate signals across nearby human cells, we constructed a human-only kNN graph (NNDescent, k = 500), symmetrized it, and column-normalized it to a transition matrix. We applied random walk with restart (RWR; restart probability α = 0.1, 100 iterations) to each model’s initial coverage vector, which preserves model-specific signal while diffusing it along the human manifold. The resulting vectors were transformed with log1p, winsorized to the 1st–99th percentiles, and min–max scaled to [0, 1] per model to obtain the final presence scores.

### Mouse Strains and Autochthonous Tumour Models

Animal studies were conducted in compliance with the ARRIVE and the European guidelines for the care and use of laboratory animals and approved by the Institutional Animal Care and Use Committees (IACUC) of the local authorities of Technische Universität München and Regierung von Oberbayern. Maximal tumour sizes of 1.5 cm or cumulative burden scores permitted by the IACUC and Regierung von Oberbayern were not exceeded in the respective studies. Animals were kept in a dedicated facility, with a light:dark cycle of 12 h:12 h, a housing temperature between 20 and 24 °C and a relative air humidity of 55%. The *LSL-Kras^G12D/+^* ^71^, *Pdx1-Cre* ^72^, *Ptf1a^Cre/+^*^73^, *Pdx1-Flp, FSF-R26^CAG-CreERT2/+^*, *FSF-Kras^G12D/+^* ^74^, *Trp53^frt^*^75^, *Trp53^lox/+^* ^76^, *Tgfbr2^flox^* ^77^, *Smad4^Lox^* ^78^, and *LSL-PIK3CA^H1047R^* ^79^ lines have been described previously. Mutant mice that develop autochthonous tumours in the pancreas were generated by intercrossing pancreas-specific Cre or Flp alleles with *LSL-Kras^G12D/+^, FSF-Kras^G12D/+^* or *LSL-PIK3CA^H1047R/+^* mice with or without floxed or FRT flanked *Trp53^frt^/Trp53^lox^, Tgfbr2^lox^*or *Smad^lox^* alleles.

### Isolation and Culture of Primary Mouse PDAC Cell Lines

Primary mouse PDAC cell lines were isolated from autochthonous pancreatic tumours and cultured in DMEM supplemented with FCS (10% v/v) and Pen/Strep (1% v/v) as previously described ^80^. All murine PDAC cell lines that were used for implantation experiments have been characterized and described previously ^81–83^. All cultures were authenticated by re-genotyping and routinely tested for mycoplasma contamination by PCR.

### Cellular Barcoding using LARRY Expressed Barcode Library

For clonal and state-fate analysis by single-cell RNA-seq, primary mouse PDAC cells were clonally tagged with expressed DNA barcodes using the LARRY Barcode Library (Addgene #140024; RRID: Addgene 140024) ^35^. Lentiviral particles were generated by co-transfecting the LARRY-EGFP library with packaging and envelope plasmids psPAX2 (Addgene, #12260) and pMD2.G (Addgene, #12259) into HEK293FT cells using the TransIT-LT1 (Mirus Bioscience) transfection reagent as previously described ^82^. Lentiviral supernatant was collected 48- and 72-hours post-transfection and sterile filtered (0.45 µm).

Primary mouse PDAC cells were transduced with lentiviral LARRY-EGFP library by spinfection (2 h, 1,000 x g, 33 °C) at a low multiplicity of infection (MOI) between 0.1 to 0.3 in 12-well plates and 1 × 10^6^ cells per well to ensure that each of the transduced cells received only one barcode. Successfully transduced-EGFP cells were sorted using a BD FACSAria Fusion Flow Cytometer and subsequently expanded.

### Orthotopic Implantation and in vivo Treatments

Orthotopic implantation was performed as previously described ^82^. In brief, mouse PDAC cells (2,500–100,000) were orthotopically implanted into the pancreas of 8-12-weeks old syngeneic immunocompetent C57Bl/6J wild-type or EGFP-tolerant Gt(ROSA)26Sor^tm1.1(CAG-cas9*,-EGFP)Fezh^/J (Cas9-EGFP) mice.

Tumour growth was monitored by magnetic resonance imaging (MRI) at two- and three-weeks post-implantation. Once tumours reached ∼100 mm³, mice were randomized into control and treatment groups. Treated groups received trametinib (3 mg kg⁻¹, daily, oral gavage), nintedanib (50 mg kg⁻¹, daily, oral gavage), or anti-PD-L1 antibody (200 µg per mouse, every third day, intraperitoneally) ^82^.

### Mouse Tumour Sample Preparation for scRNA-seq

Mouse PDAC samples for scRNA-seq were prepared as previously described ^82^. Tumours were dissociated and enzymatically digested using the Tumor Dissociation Kit (Miltenyi Biotec, #130-096-730) for 40 min at 37 °C with agitation. The resulting cell suspension was filtered through a 100-µm strainer, pelleted, and resuspended in 2% FCS/PBS containing RNase inhibitor (New England Biolabs, # M0314L, 1:1000). Cell debris was removed using Debris removal solution (Miltenyi, catalog #130-109-398) and live cells were enriched using the Dead Cell Removal Kit (Miltenyi Biotec, #130-090-101), both according to the manufacturer’s instructions. Cells were resuspended in PBS and blocked for 10 min on ice with anti-mouse TruStain FcX™ (anti-mouse CD16/32, BioLegend, 1:100) to prevent nonspecific antibody binding. If cell subset enrichment was performed prior to scRNA-sequencing, suspensions were subjected to fluorescence-activated cell sorting (FACS) or magnetic-activated cell sorting (MACS). For FACS, cells were stained with Ter-119-BV421 (BioLegend, 1:100), CD45-AF647 (BioLegend, 1:20), CD31-AF647 (BioLegend, 1:20), and EPCAM-AF647 (BioLegend, 1:20) for 30 min on ice and sorted using a BD FACSAria Fusion. The TER-119-/CD45+/CD31+/EPCAM+ fraction (enriched for immune, endothelial, and epithelial tumour cells, but exclusion of erythrocytes) and the TER-119-/CD45-/CD31-/EPCAM- fraction (enriched for fibroblasts/mesenchymal tumour cells, and exclusion of erythrocytes) were collected in 2% FCS/PBS. For MACS enrichment, cells were incubated with CD45 MicroBeads (Miltenyi Biotec, #130-052-301) and separated on MS columns according to the manufacturer’s instructions. Both the positively selected CD45⁺ cells and the flow-through (enriched for non-immune cells) were collected in 2% FCS/PBS for downstream library preparation.

### Flow Cytometry Staining of Mouse Tumour Samples

Up to two million cells from dissociated tumour suspensions were stained in 96-well V-bottom plates. First, cells were washed with PBS and stained with fixable live/dead dye iFluor^TM^840 maleimide (AAT Bioquest) and anti-mouse TruStain FcX™ (anti-mouse CD16/32, BioLegend, 1:100), to block Fc receptors (15 min, 4°C). After washing with FACS buffer (PBS + 0.5% BSA, 0.1% NaN_3_), extracellular staining was performed with marker antibodies for 30 min at 4 °C. Biotinylated antibodies was applied using a two-step staining protocol. Cells were then fixed with the eBioscience™ Foxp3/Transcription Factor Staining Buffer Set (Thermo Fisher Scientific, #00-5523-00) for 30 min at RT, washed twice with permeabilization buffer, and stored in FACS buffer at 4 °C until acquisition. The samples were acquired on a CytoFLEX LX flow cytometer, gating on single, viable CD45⁺ or CD45⁺ CD3E⁺ cells. The following antibodies were used: CD45-BUV737 (BD, 1:200), CD11b-eF605NC (eBioscience, 1:100), CD3e-biotin (BD, 1:200), CD4-BUV496 (BD, 1:200), CD8a-BUV785 (BioLegend, 1:200), and Streptavidin-BUV395 (BD, 1:200).

### scRNA-seq Library Preparation and Sequencing

Single-cell GEM generation, barcoding, and library construction were performed using the 10x Genomics Chromium Single Cell 3’ Reagent Kits (v3.1 Chemistry) according to the manufacturer’s instructions. Cells were counted, diluted in 2% FCS/PBS, and up to 35,000 cells were loaded per lane on a Chromium Next GEM Chip G to generate gel beads in emulsion. cDNA and libraries were assessed for quality and fragment size using an Agilent Bioanalyzer 2100 System with the High Sensitivity DNA Kit (Agilent, #5067-4626). Libraries were sequenced on an Illumina NovaSeq 6000 S2 flow cell with paired end reads.

## Code and data availability

The human and publicly available mouse datasets can be downloaded from public sources (Supplementary Table 1-2). The processed Atlases, the raw in-house mouse datasets, and the code will be made public upon publication.

## Acknowledgments

We thank Aviv Regev and Malte Lücken and their respective laboratories members for their insightful discussions, valuable feedback, and generous sharing of expertise and resources that greatly contributed to this work. This manuscript used data generated by the TCGA Research Network. This study was supported by the German Cancer Consortium (DKTK) and DKTK Joint Funding, the DKFZ-MOST cooperation program (Ca-217), the European Research Council (Consolidator grant CoG PACA-MET-819642 and MSCA ITN-ETN-861196 to R.R.; CoG - 648521 to D.S.; Advanced Grant AdG DeepCell – 101054957 to F.T.); the Deutsche Forschungsgemeinschaft (SFB 1371 Project-ID 395357507 P12 to D.S.; DFG SA 1374/8-1 Project-ID 515991405 to D.S.; DFG SA 1374/7-1 Project-ID 515571394 to D.S., M.S.S., F.T.; DFG SA 1374/6-1 Project-ID 458890590 to D.S.); Deutsche Krebshilfe (#70117118 DEFEAT-PDAC of the German Pancreatic Cancer Alliance to D.S., R.R., M.S.S., G.S., F.T., C.F.; #70115743 to D.S.; #70116843 to D.S.); the German Federal Ministry of Education and Research (DROP2AI #031L0305B to D.S., G.S., R.R.), the Wilhelm Sander-Stiftung (2020.174.1 and 2017.091.2 to D.S.), and the Chan Zuckerberg Initiative Foundation (CZIF; grant CZIF2022-007488 (Human Cell Atlas Data Ecosystem to F.T.). C.F. is a Cancer Research Institute Irvington Fellow supported by the Cancer Research Institute (CRI4641).

## Author contributions

D.L. and D.S. conceived the study. D.L. and S.P. contributed equally and have the right to list their name first in their curriculum vitae. D.L., and S.P. led, conceptualized, and organized the project with support from S.J. D.L. and S.P. led the analysis of the atlas, with support from P.P., J.M.A.-Q. and R.F.K. Data curation was performed by D.G. C.S., D.A.N., P.P., T.K., D.W., V.G., M.K., C.S., M.R.L. and T.B.H. D.A.N., M.Z., T.K., M.Z., B.S., C.F. and S.B. generated the in-house mouse datasets. F.J.T. and D.S. directed and supervised the work with support from S.J., G.S., R.R., C.F., M.S.S., L.H., and WLH. D.S. and F.J.T. acquired funding. All authors contributed to writing, reviewing and editing the manuscript, and approved the final version. Y.D., and M.K. contributed to the figure design.

## Extended data figures and tables

Extended Data Fig. 1–5 and Supplementary Tables 1-10.

