## Extended figures for "Cross-species single-cell atlases chart progression, therapy-driven remodelling and immune evasion in pancreatic cancer"

### Extended Data Figures and legends

**Extended Figure 1**

**a**

| Method | Bio conservation: Isolated labels | Bio conservation: KMeans NMI | Bio conservation: KMeans ARI | Bio conservation: Silhouette label | Bio conservation: cLISI | Batch correction: Silhouette batch | Batch correction: iLISI | Batch correction: KBET | Batch correction: Graph connectivity | Batch correction: PCR comparison | Aggregate score: Batch correction | Aggregate score: Bio conservation | Aggregate score: Total |
|---|---|---|---|---|---|---|---|---|---|---|---|---|---|
| **scanvi_emb** | 0.71 | 1.00 | 1.00 | 1.00 | 0.81 | 0.03 | 0.93 | 0.34 | 0.53 | 0.47 | 0.46 | 0.90 | 0.73 |
| **expimap_emb** | 0.67 | 0.64 | 0.44 | 0.46 | 0.70 | 0.79 | 0.93 | 1.00 | 0.61 | 1.00 | 0.87 | 0.58 | 0.70 |
| **drvi_emb** | 0.29 | 0.81 | 0.98 | 0.90 | 0.81 | 1.00 | 0.75 | 0.42 | 0.00 | 0.48 | 0.53 | 0.76 | 0.67 |
| **scpoli_emb** | 0.51 | 0.75 | 0.36 | 0.80 | 0.80 | 0.11 | 0.77 | 0.88 | 1.00 | 0.38 | 0.63 | 0.64 | 0.64 |
| **sysvi_emb** | 0.00 | 0.23 | 0.15 | 0.16 | 0.51 | 0.23 | 0.74 | 0.09 | 0.78 | 0.78 | 0.53 | 0.21 | 0.34 |
| **scvi_emb** | 1.00 | 0.00 | 0.00 | 0.35 | 0.00 | 0.00 | 1.00 | 0.14 | 0.43 | 0.60 | 0.43 | 0.27 | 0.34 |
| **X_pca** | 0.23 | 0.35 | 0.44 | 0.00 | 1.00 | 0.68 | 0.00 | 0.00 | 0.25 | 0.00 | 0.19 | 0.41 | 0.32 |

**b**

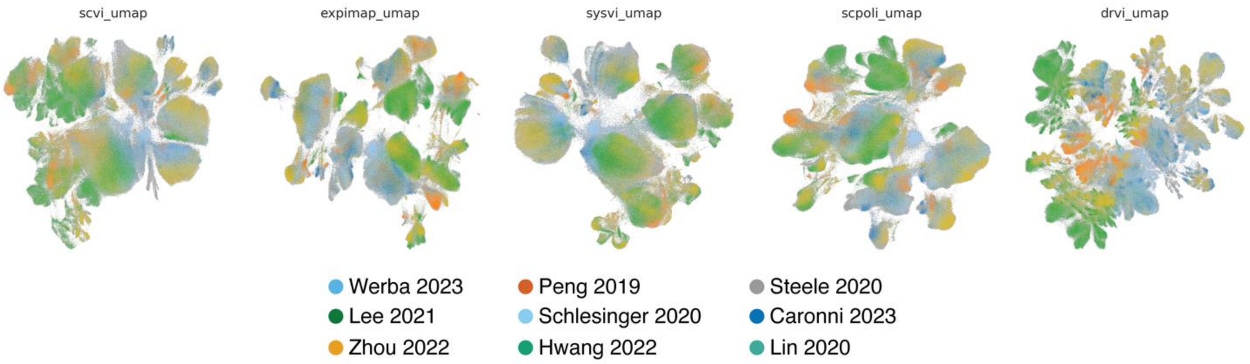


**Extended Data Fig. 1. Integration methods for batch correction. a)** ScIB Metrics show scANVI has the best batch correction performance. **b)** UMAP embeddings derived from the other benchmarked batch correction methods.

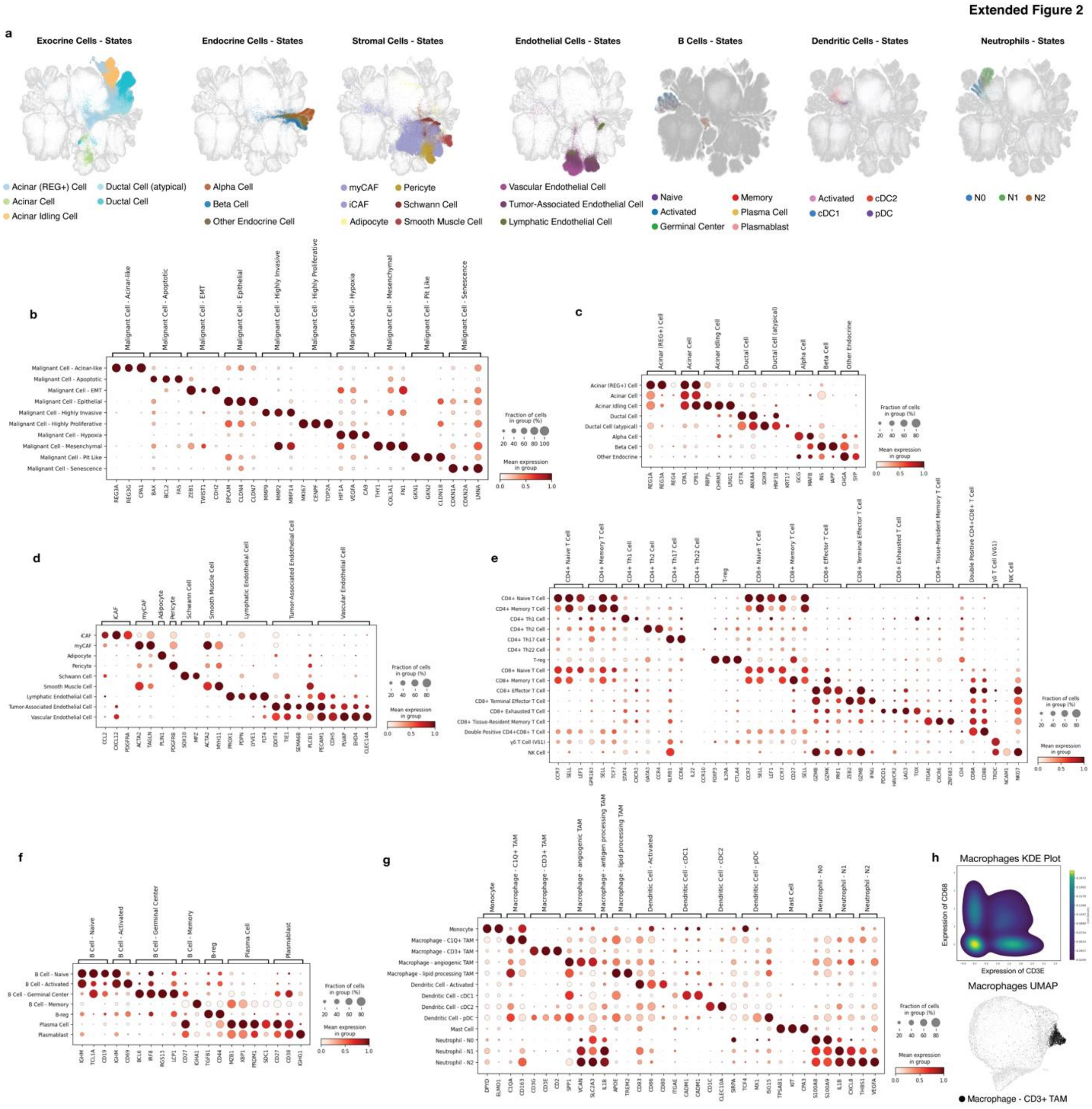


**Extended Data Fig. 2. Additional human PDAC cell states identified from patient transcriptomes. a)** UMAP embedding of exocrine, endocrine, stromal, endothelial, B cells, dendritic cells and neutrophils highlighting cell states. **b-g)** Dotplots of marker genes used to support state-level annotations across major compartments, including malignant cells (b), exocrine and endocrine cells (c) stromal (pericyte/schwann/smooth muscle/CAF (myCAF/iCAF), and endothelial cells) (d), T cells (e), B cells (f), and myeloid cells (g). **h)** KDE density plot of macrophages expressing CD68 and CD3E highlighting the rare cell type presence (top). UMAP of all macrophages in the human atlas with the CD3+ TAM highlighted in black (bottom).

**Extended Figure 3**

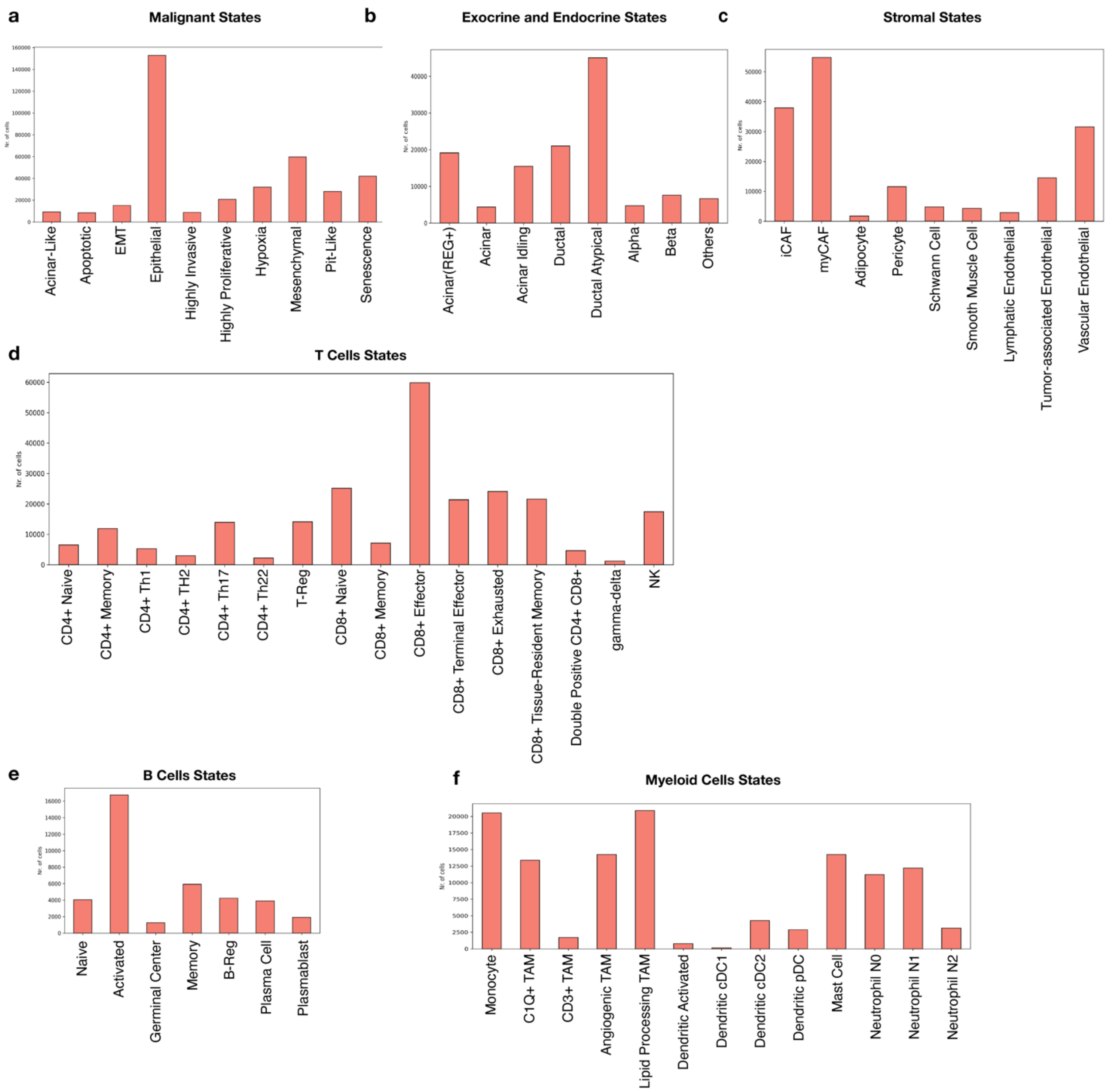


**Extended Data Fig. 3. Cell numbers of level 4 annotations of the Human PDAC atlas. a-f)** Absolute number of cells for each category in the level 4 annotation.

Extended Figure 4

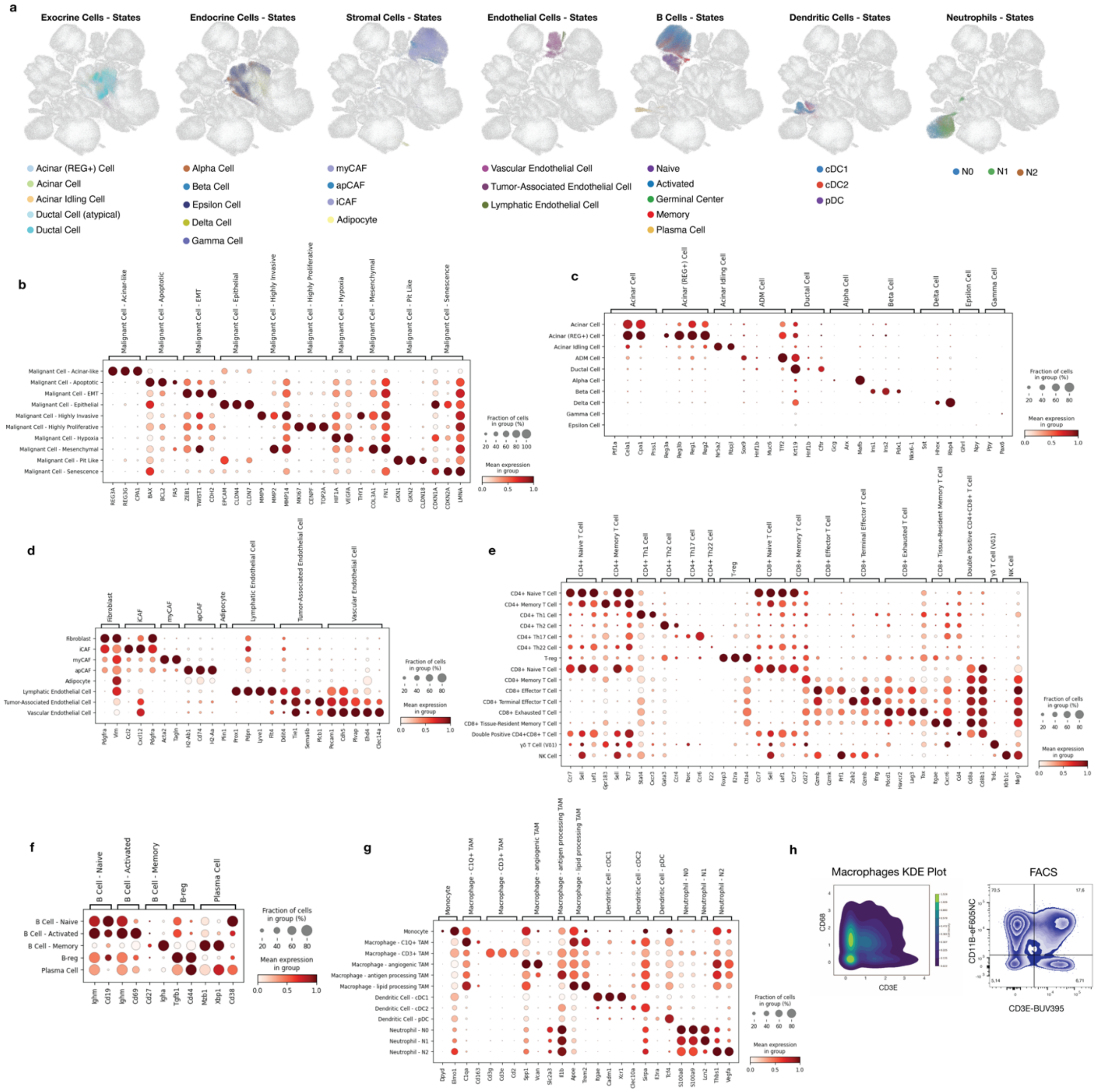


**Extended Data Fig. 4. Additional mouse PDAC cell states identified from mouse models. a)** UMAP embedding of Exocrine, Endocrine, Stromal, Endothelial, B cells, Dendritic cells and Neutrophils highlighting cell states. **b-g)** Dotplots of marker genes used to support state-level annotations across major compartments, including malignant cells (b), exocrine and endocrine cells (c) stromal, adipocyte/CAF (myCAF/iCAF/apCAF) and endothelial cells (d), T cells (e), B cells (f), and myeloid cells (g). **h)** Validation of macrophage subsets using transcriptional density plots of markers (CD68/CD3E) and protein-based flow-cytometry (FACS) derived density plots (CD11/CD3).

**Extended Figure 5**

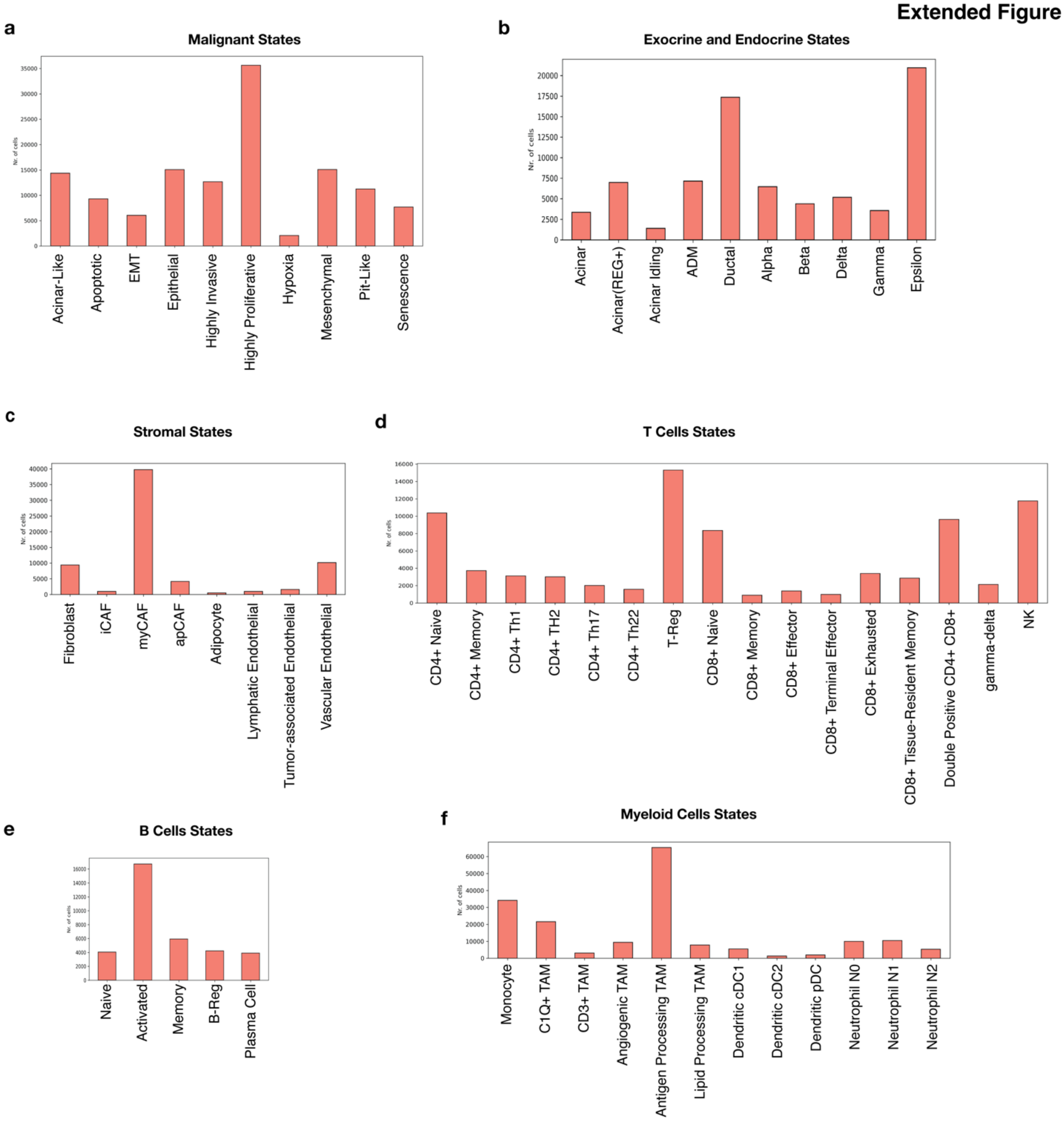


**Extended Data Fig. 5. Cell numbers of level 4 annotations of the Mouse PDAC atlas. a-f)** Absolute number of cells for each category of the mouse PDAC atlas in the level 4 annotation.
